# Deformability screening identifies NUDT5 as a mediator of cellular mechanobiology

**DOI:** 10.64898/2026.07.15.734864

**Authors:** Angelina M. Flores, Jennifer Soto, Navjot Kaur Gill, Ahmed Almunaifi, Chau Ly, Dongping Qi, Ruchira Krishnamurthy, Eleana Parajón, Bobby Tofig, Valeria Garcia, Maria Sol Recouvreux, Song Li, Yi Lu, Beth Y. Karlan, Douglas N. Robinson, Junyoung O. Park, Robert Damoiseaux, Sandra Orsulic, Amy C. Rowat

## Abstract

How cells deform, sense, and respond to mechanical cues drives physiological and disease processes ranging from development to cancer metastasis; however, unbiased approaches to identify mechanical mediators are lacking. We screened 1280 compounds to identify modulators of cancer cell deformability using a cellular filtration assay and identified 92 compounds that significantly reduced deformability of ovarian cancer cells; top hits also reduced migration and invasion. Connectivity mapping of the top 21 compounds identified NUDT5 (Nudix hydrolase 5) as a predicted mechanical mediator; transcriptomic analyses implicated NUDT5 in mechanobiology and metabolic processes. We confirmed that NUDT5 mediates intracellular ATP and cellular mechanical behaviors, including morphology and deformability. In ovarian cancer, increased NUDT5 levels were associated with higher tumor stage and worse patient survival; NUDT5 inhibition reduced migration and colony formation *in vitro* and peritoneal tumor burden in mice. These findings establish deformability-based screening as a platform for discovering mechanical mediators and identify NUDT5 as a therapeutic target in ovarian cancer.

**Teaser:** Screening cells based on deformability provides an unbiased approach to identify NUDT5 as a mediator of cell mechanics

## INTRODUCTION

Cellular mechanobiology—how cells deform, sense, and respond to mechanical cues— underlies fundamental processes from tissue morphogenesis (*1*) to wound healing (*2*) to cancer metastasis (*3*). For example, the increased cellular deformability of cancer cells is associated with mesenchymal and amoeboid phenotypes (*4, 5*), enhanced motility and invasion (*6*), and resistance to chemotherapy drugs (*7, 8*). Mapping the molecular mediators that govern cellular mechanical behaviors such as cell shape, cell stiffness or deformability, and motility, would enable predictive insights and deliberate control of cells to block disease-relevant phenotypes, such as the cellular shape changes required for confined migration and invasion. Identifying modulators of cellular mechanical behaviors could further reveal functional biomarkers of disease progression and potential therapeutic targets. However, the molecular mediators underlying cell mechanical behaviors remain incompletely understood, and existing methods lack the throughput needed for unbiased discovery.

To establish molecular mediators of cellular mechanical behaviors—or mechanical mediators—studies of cell mechanics have historically focused on individual proteins including cytoskeletal proteins (*9, 10*) and nuclear lamins (*11*), and signaling axes such as RhoA/ROCK (*12*). Such studies have typically used low-throughput approaches that involve applying mechanical force to individual cells sequentially using atomic force microscopy (AFM) (*13*) or optical stretching (*14*). While these methods have advanced mechanistic knowledge of cell mechanobiology, they require substantial user input and computationally intensive analysis, thereby hindering larger-scale screening applications. Recent developments have increased throughput for quantifying cellular rheology (*15*) and traction stresses (*16*). However, methods for measuring cellular deformability remain incompatible with existing high throughput screening (HTS) platforms, and screening >10³ samples based on cellular deformability to enable the systematic discovery of mechanical mediators remains inaccessible.

Here, we scaled our cellular filtration assay to screen 1,280 small molecules of the Library of Pharmacologically Active Compounds (LOPAC) for modulators of deformability in cisplatin-resistant human ovarian cancer (OVCAR-5-CisR) cells, which are more deformable than drug-sensitive cells (*7*). The filtration approach requires cells to rapidly transit through micron-scale pores in response to applied pressure on timescales of seconds; deformable cells can readily pass through, whereas stiffer cells occlude pores and reduce filtration (*7*). Integrating deformability screening with transcriptomic connectivity mapping, we identify NUDT5, an enzyme that regulates intracellular ATP, as a mechanical mediator that controls cellular stiffness, morphology, and migration and is associated with ovarian cancer progression.

## RESULTS

### Deformability screen identifies small molecules that modulate cell mechanical behaviors

To screen cells based on deformability, we applied air pressure to drive a suspension of cells to filter through a porous membrane (10 µm pores) over timescales of seconds and measured the retained volume, or ‘retention’ (**Fig. 1A**) (*7*); this served as a readout of deformability. A lower retention volume indicates cells that readily deform and transit through the pores, whereas a higher retention volume indicates pore occlusion and reduced deformability. The short filtration timescale of seconds captures the intrinsic deformability of cells, rather than migration.

**Fig. 1.**
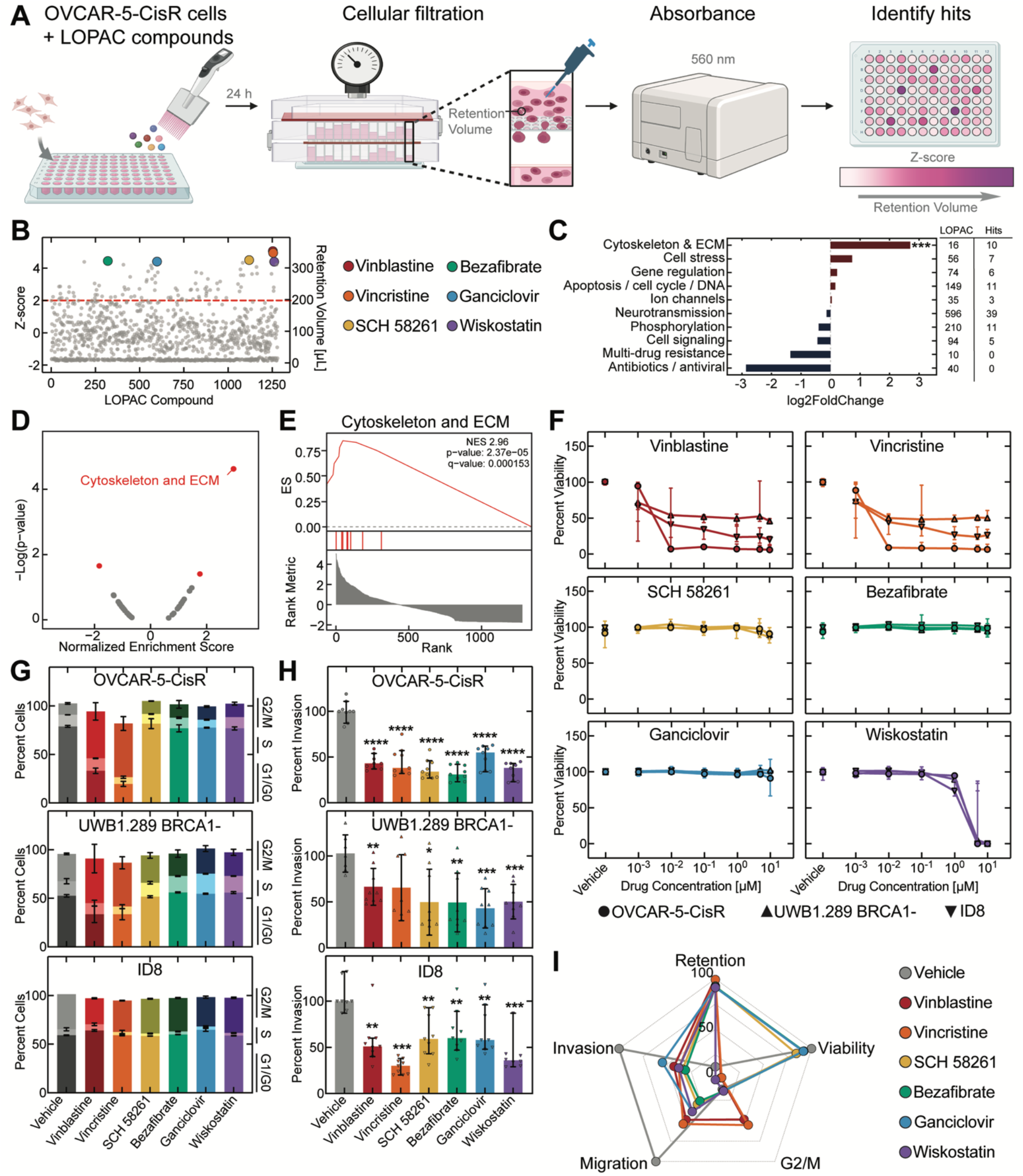
Identifying small molecules that impact cell deformability. (**A**) Workflow of the deformability screen. Human ovarian cancer (OVCAR-5-CisR) cells were treated with the 1280 compounds of the LOPAC library and subject to pressure-driven filtration through 10 µm pores. Retention volume was measured by absorbance (560 nm) to identify top compounds. Created with Biorender.com. (**B**) The top six hits: vinblastine (red); vincristine (orange); SCH 58261 (yellow); bezafibrate (green); ganciclovir (blue); and wiskostatin (purple); 92 compounds had Z-score > 2 (dashed red line). (**C**) Log_2_ fold enrichment of drug categories in the top 92 compounds relative to the LOPAC library. ***p < 0.0001. (**D**) Drug set enrichment analysis of LOPAC compounds ranked by Z-score showing normalized enrichment score by drug category. (**E**) Enrichment scores (ES) for cytoskeleton and ECM drug class. (**F**) Cell viability by luminescence assay (48 h drug treatment). Human ovarian cancer cell lines: OVCAR-5-CisR (circle); UWB.289 BRCA1-(upward triangle); or murine ID8 (downward triangle) cells. (**G**) Cell cycle distribution by flow cytometry (48 h drug treatment). (**H**) Transwell invasion assay through basement membrane extract (BME)-coated membranes (8 µm pores) at 24 h (24 h drug treatment). Data shows median with 95% confidence interval for **F** - **H**. (**I**) Radar plots summarize OVCAR-5-CisR results from **B and F** (10 µM)**, and** from **G-H** and **S1D**: 0.5 nM for vinblastine and vincristine; 0.1 µM for wiskostatin; and 1 µM for SCH 58261, bezafibrate, and ganciclovir. For all data n = 3 independent experiments, except n = 1 for **B**, and n = 2 for **G**.

As a model, we used cisplatin-resistant OVCAR-5-CisR cells, which are more deformable than parental OVCAR-5 cells, consistent with their mesenchymal phenotype (*7*). We previously showed that paclitaxel – which stabilizes microtubules and stiffens cells – increases retention in a concentration-dependent manner (*7*). We therefore used paclitaxel as a positive control to optimize assay conditions, targeting Z’ ≥ 0.5, as the accepted standard for high-throughput screening assay validation (*17*); the subsequent screen was performed under these validated conditions. Since cellular deformability and motility share common mediators (*18, 19*), we hypothesized that top compounds identified in the screen to reduce cell deformability would also impact disease-relevant cell behaviors, such as invasion and migration.

We screened all 1,280 compounds of the LOPAC collection, which span diverse mechanisms of action (**fig. S1A**); this enabled systematic interrogation of pathways that modulate cellular deformability. To quantify retention volume across each 96-well plate, we measured the absorbance of phenol red (560 nm) in cell culture media using a plate reader. To prioritize small molecules that have the most significant effects on cell deformability compared to vehicle (DMSO)-treated cells, we ranked the compounds based on their Z-score, *Z = (Absorbance_Drug_ - Mean Absorbance_Vehicle_) / SD Absorbance_Vehicle_* ; this revealed 92 compounds with significantly increased retention volume (Z > 2) (**Fig. 1B and Table S1**). Among these 92 compounds, we found a significant enrichment in compounds targeting pathways involved in ‘cytoskeletal and extracellular matrix (ECM)’ (log_2_ fold change, log_2_FC = 2.7; p = 1.0 × 10^-9^) (**Fig. 1C-E**), which represents a ∼9-fold increase in prevalence compared to the original LOPAC library. By contrast, there was no enrichment among top hits for compounds in the neurotransmission category despite the LOPAC library consisting of 47% of compounds in this category (log_2_FC = -0.16; p = 0.73) (**Fig. 1C, fig. S1A**). The top two compounds, vinblastine and vincristine, are well-established regulators of microtubule dynamics (*20, 21*). We also identified wiskostatin, which inhibits the Arp-2/3 complex that regulates cellular motility and stiffness (*18*). These findings support that deformability-based screening preferentially identifies compounds that modulate cell mechanics.

To characterize the top six hits (Z-score > 4.4; 0.5% of library) – vinblastine, vincristine, SCH 58261, bezafibrate, ganciclovir, and wiskostatin – we first assessed cell viability across three cell lines (human ovarian cancer OVCAR-5-CisR, UWB1.289 BRCA1-, and murine ID8 cells; **Fig. 1F**). In cytotoxicity assays, vinblastine and vincristine markedly reduced cell viability at drug concentrations above 0.01 µM for OVCAR-5-CisR, and above 0.001 µM for UWB1.289 BRCA1- and ID8 cells. With wiskostatin treatment, all cells remain viable up to 10 µM. By contrast, SCH 58261, bezafibrate, and ganciclovir showed minimal to no changes in cell viability across 0.001 to 10 µM (**Fig. 1F, fig. S1B**), supporting that the increased retention can occur independently of cytotoxic effects. Guided by these concentration-response data, we selected the following sublethal doses of the lead compounds for subsequent assays with OVCAR-5-CisR, UWB1.289 BRCA1-, and ID8 cells: 0.5 nM for vinblastine and vincristine; 0.1 µM for wiskostatin; and 1 µM for SCH 58261, bezafibrate, and ganciclovir.

As cell cycle stage can impact cellular deformability (*22*), we tested for drug-induced changes in cell cycle distribution using flow cytometry of propidium iodide-labeled cell populations (**fig. S1C**). While vinblastine and vincristine had no effects on cell cycle progression in ID8 cells, these drugs induced G2/M arrest in OVCAR-5-CisR and UWB1.289 BRCA1-cells (**Fig. 1G**), consistent with their known effects on microtubule dynamics (*20, 21*). By contrast, SCH 58261, bezafibrate, ganciclovir, and wiskostatin did not significantly alter cell cycle distribution in any of the cell lines tested (**Fig. 1G**). These findings indicate that while some top compounds alter cell cycle distribution, the screen also reveals compounds that alter deformability independently of cell cycle changes.

Cell deformability and motility are regulated by shared molecular mediators (*18*), so we next assessed cell invasion, one of the hallmark features of cancer cells that reflects their ability to undergo the large shape changes required to migrate through confined spaces, invade the basement membrane, and colonize healthy tissues. To measure cell invasion, we quantified the number of cells that invade through 8 µm pores coated with a basement membrane extract (BME) matrix using a transwell assay (**Fig. 1H**). We also measured the migration of cells through the 8 µm pores of uncoated transwell membranes (**fig. S1D**). We found that all top six compounds consistently reduced cell invasion and migration at sublethal concentrations.

To visualize trends of how the top six compounds impact cellular phenotypes across assays, we generated a radar plot that summarizes findings for OVCAR-5-CisR cells as shown in **Fig. 1I**. The top two lead compounds, vinblastine and vincristine, showed marked effects on cellular deformability (increased retention volume) from the deformability screen; significantly reduced cell migration and invasion; blocked cell cycle progression; and at higher concentrations had cytotoxic effects. The other top four compounds – SCH 58261, bezafibrate, ganciclovir, and wiskostatin – significantly reduced deformability, as shown by the increased retention volume, and reduced migration and invasion but otherwise did not significantly impact viability or cell cycle regulation (**Fig. 1F-I**). Together, these findings show that deformability screening can identify small molecules that also reduce cell migration and invasion.

### NUDT5 as a predicted mechanical mediator

We next investigated if there were common molecular mediators that contributed to the reduction in cellular deformability induced by the mechano-modulating drugs that emerged from the screen. To identify shared mediators of cellular deformability, we performed a transcriptomic connectivity analysis using the publicly accessible Connectivity Map (cMAP) database (*23*). We queried the top 35 compounds (Z-score > 3), of which 21 had available drug-induced gene expression data; genetic knockdown (KD) signatures that mirror the drug-induced gene signatures associated with the reduced cell deformability are indicated by a positive connectivity score (**Fig. 2A**). This meta-analysis revealed established mediators of cell mechanical properties including the GTPase H-Ras (HRAS) (*7, 24*), focal adhesion kinase (PTK2) (*25, 26*), and epidermal growth factor receptor (EGFR) (*27*). Many of these genes are also implicated in epithelial-to-mesenchymal transition (EMT) and cancer progression (*28–30*). Notably, we identified NUDT5, a member of the Nudix hydrolase family that catalyzes the formation of ATP from ADP-ribose to regulate chromatin remodeling and DNA damage repair (*31, 32*). NUDT5 also plays a non-enzymatic role in nucleotide homeostasis (*33–35*). While NUDT5 has been studied in breast cancer (*36–38*), its roles in cell mechanical regulation and ovarian cancer remain unknown.

**Fig. 2.**
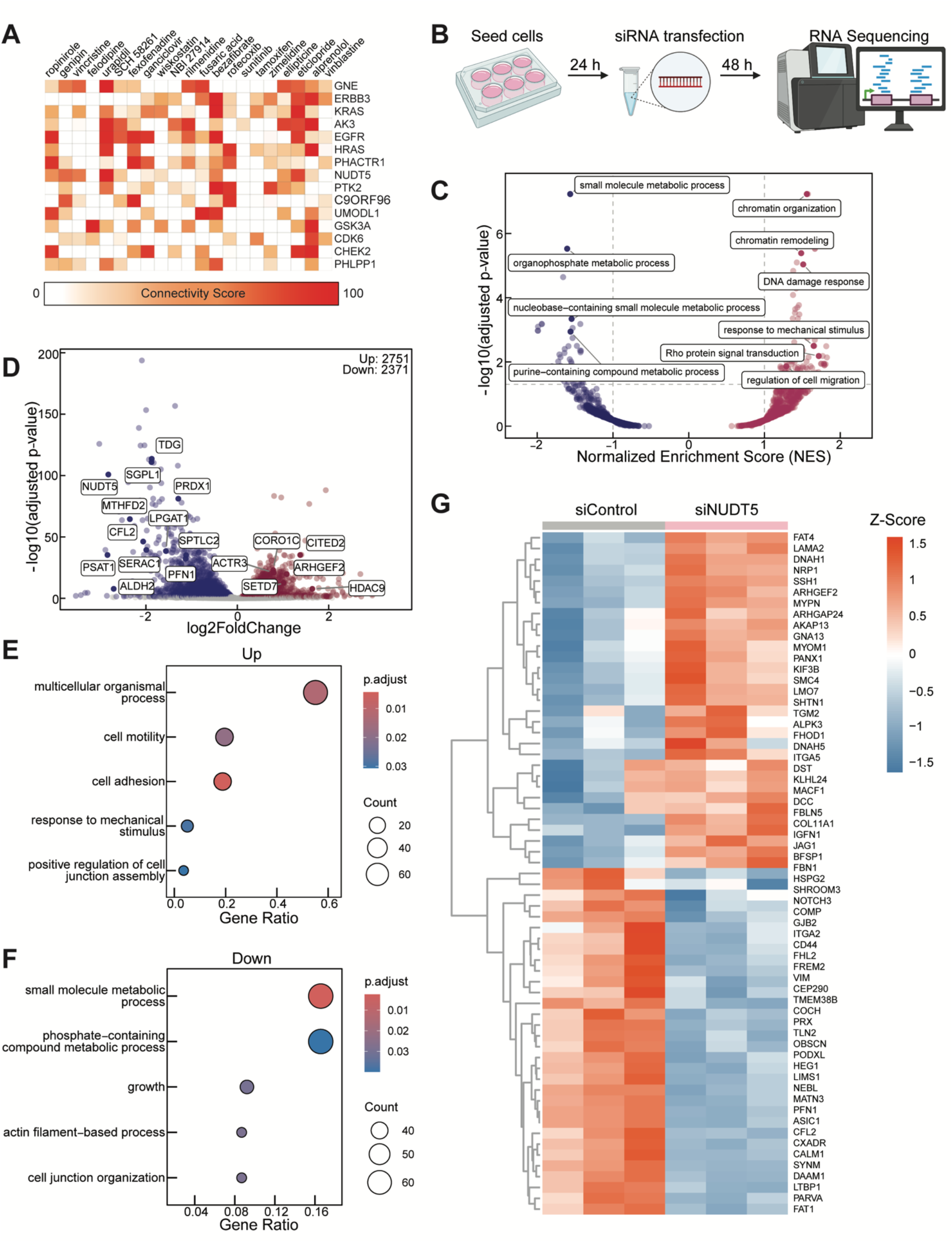
NUDT5 as a predicted mechanical mediator. (**A**) Transcriptomic connectivity analysis of 21 of the top compounds (Z > 3) from the deformability screen. Shown here are the top 15 genes with the highest combined connectivity scores across 8 different epithelial cell lines. (**B**) Schematic of RNA-sequencing (RNAseq) workflow. Created with Biorender.com. (**C**) Volcano plot of enriched Gene Ontology (GO) terms from RNAseq data of siNUDT5 and siControl OVCAR-5-CisR cells at day 2 post-transfection. (**D**) Differentially expressed genes (DEGs) in siNUDT5 versus siControl: upregulated DEGs (red, p_adj_ < 0.05), downregulated DEGs (blue, p_adj_ < 0.05); non-significant DEGs (grey, p_adj_ > 0.05). (**E, F**) Overrepresentation analysis of upregulated (**E;** log_2_FC > 1, p_adj_ < 0.05) and downregulated (**F;** log_2_FC < -1, p_adj_ < 0.05) DEGs. Significantly altered GO terms are ranked by gene ratio (number of genes per GO term divided by the total number of significant genes). Colors represent the adjusted p-values (p_adj_), and dot size reflects the gene count. (**G**) Heatmap of DEGs (log_2_FC > 0.5, p_adj_ < 0.01) that are established mechanical mediators as defined as Mechanobiology Priority Score ≥ 80 (*40*). Color represents changes in mRNA levels based on Z-score. RNASeq data from OVCAR-5-CisR cells represent the average of n = 3 independent experiments.

To validate NUDT5 as a predicted mechanical mediator, we first measured the expression of NUDT5 in two established ovarian cancer cell lines, OVCAR-5-CisR and PEA-1. We selected PEA-1 cells because they showed the highest NUDT5 levels among ovarian cancer cells in the Cancer Cell Line Encyclopedia (CCLE) database (**fig. S2A**). We quantified levels of the three distinct isoforms of NUDT5 in OVCAR-5-CisR and PEA-1 cells using qRT-PCR; as shown in **fig. S2B**, NUDT5 isoform 1 was predominant in both cells, with ∼70× higher transcript levels than isoforms 2 and 3. PEA-1 cells showed higher levels of NUDT5 compared to OVCAR-5-CisR cells at both transcript and protein levels (**fig. S2B-C**). We next assessed the efficacy of siRNA mediated knockdown of NUDT5 (siNUDT5) in OVCAR-5-CisR cells, confirming that siNUDT5 reduced mRNA levels of NUDT5 isoforms 1 and 2 by >80% (**fig. S3A**). Consistent with these findings, total protein was reduced by >80% for up to 5 days in OVCAR-5-CisR and over 3 days in PEA-1 cells as determined by immunoblotting (**fig. S3B-G**). Immunostaining further confirmed that siNUDT5 reduced levels of NUDT5 in both cytoplasmic and nuclear regions (**fig. S3H-I**).

Having established knockdown of NUDT5 in ovarian cancer cells, we characterized the transcriptional effects of NUDT5 depletion in OVCAR-5-CisR cells using RNA-sequencing (**Fig. 2B**). Principal component analysis revealed distinct clustering of siNUDT5 and siControl cells (**fig. S4A**). Analysis of the differentially expressed genes (DEGs) identified a total of 2751 upregulated genes and 2371 downregulated genes in siNUDT5 cells compared to siControl cells (p-adjusted, p_adj_ < 0.05) (**Fig. 2D**). Among the top DEGs, we confirmed a significant 87% reduction of NUDT5 in siNUDT5 cells (**fig. S4B**), consistent with our qRT-PCR and immunoblotting results (**fig. S3A-E**). Since NUDT5 is a member of the NUDT superfamily, we confirmed that siNUDT5 specifically reduced NUDT5 expression; most other family members were unaffected, with modest upregulation of NUDT6, NUDT12, and NUDT22 (**fig. S4B**).

To investigate how NUDT5 impacts biological processes, we conducted gene set enrichment analysis; this revealed significant enrichment in gene sets associated with chromatin remodeling, DNA damage response, as well as nucleotide and purine metabolism (**Fig. 2C**). These themes are represented among the top DEGs including SETD7, CITED2, and HDAC9 (chromatin remodeling); TDG and PRDX1 (DNA damage response); and MTHFD2 and PSAT1 (purine metabolism) (**Fig. 2D**). Overrepresentation analyses of the downregulated DEGs further confirmed the effects of NUDT5 depletion on small molecule and phosphate-containing metabolic processes **(Fig. 2F)**. These findings are consistent with previous reports of the enzymatic and non-enzymatic roles of NUDT5 in mediating ATP and adenosine metabolite levels for DNA damage repair and nucleotide homeostasis (*31–35, 39*).

Interestingly, depletion of NUDT5 also significantly altered the expression of genes implicated in cellular mechanobiology, with an enrichment of gene sets related to response to mechanical stimulus, Rho protein signal transduction, and cell migration (**Fig. 2C**). Overrepresentation analyses further showed that the top upregulated DEGs (p_adj_ < 0.05, log_2_FC > 1.0) were associated with cell adhesion and motility, while top downregulated DEGs (p_adj_ < 0.05, log_2_FC < -1.0) were associated with actin-filament based processes (**Fig. 2E-F**). Indeed, top DEGs include the actin-regulatory genes profilin 1 (PFN1), cofilin 2 (CFL2), coronin 1C (CORO1C), and actin related protein 2/3 (ACTR3), which is a core component of the Arp2/3 complex. Analysis of the DEGs in the context of established mechanical mediators (*40*) identified additional differentially expressed mechano-regulatory genes that mediate cytoskeletal structure and mechanics (the intermediate filament proteins, vimentin, VIM, and synemin, SYNM); cell motility (Rho GTPase Activating Protein 24, ARHGAP24, and Rho/Rac Guanine Nucleotide Exchange Factor 2, ARHGEF2); and cell-matrix interactions (integrins ITGA5, ITGA2 and the ECM protein fibrillin 1, FBN1) (**Fig. 2G**). These findings suggest a previously unrecognized role for NUDT5 in modulating cell mechanical behaviors.

### NUDT5 modulates cell mechanical behaviors and ATP production

We next evaluated how NUDT5 regulates cellular mechanical behaviors using a series of *in vitro* assays. We first tested cell deformability using the filtration assay (as in **Fig. 1A**). Cells with NUDT5 depletion were more deformable than controls, with decreased retention across both OVCAR-5-CisR and PEA-1 cells (**Fig. 3A-B**). While there is some variability across the two siRNA oligonucleotides against NUDT5, the trend of decreased retention with NUDT5 depletion is similar to the effects of pharmacologic perturbation of F-actin using cytochalasin D, which has been well-characterized to increase cell deformability (*41*). We confirmed that siNUDT5 cells have a similar cell size distribution as siControl cells (**fig. S5A-B**), indicating that altered cell mechanics rather than a reduction in cell size determines the increased filtration.

**Fig. 3.**
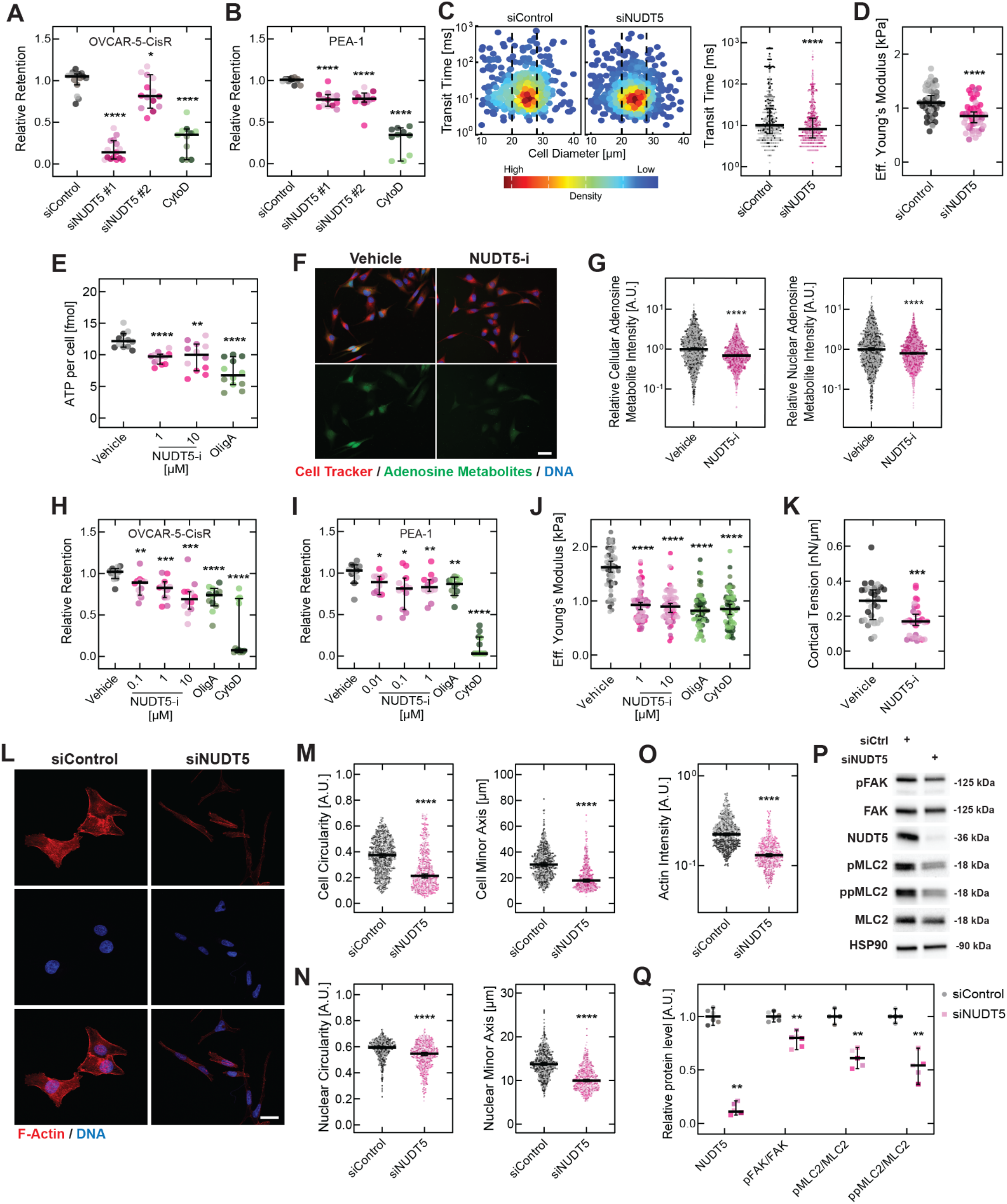
NUDT5 modulates cell mechanical behaviors and ATP production. (**A, B**) Cell filtration of OVCAR-5-CisR and PEA-1 cells with siNUDT5 knockdown versus siControl (10 µm pores). Cytochalasin D (CytoD; 1 µM, 30 min) serves as a control (N ≥ 10 technical replicates). (**C**) Quantitative deformability cytometry (q-DC): transit time measures cell passage through a 9 × 10 μm constriction under applied pressure. Dashed lines indicate size-gating thresholds (N > 200 cells per condition). (**D**) Force probe indentation of adhered cells (N = 45 cells per condition). (**E**) ATP levels by luminescence assay; cells treated with vehicle (DMSO), NUDT5-i (TH5427)- or oligomycin A (OligA; 1 µM) for 24 h. (**F**) Epifluorescence images of adenosine-responsive aptamer (green), CellTracker (red), and DAPI (blue). Scale, 50 µm. (**G**) Adenosine metabolite levels in whole cell and nuclear regions (N > 1000 cells per condition). (**H, I**) Cell filtration of OVCAR-5-CisR cells and PEA-1 cells (N ≥ 11 technical replicates per condition). (**J**) Force probe indentation (N > 58 cells per condition). (**K**) Cortical tension by micropipette aspiration; vehicle or 1 µM NUDT5-i (N ≥ 29 cells per condition). (**L**) Confocal images of F-actin (phalloidin, red) and DNA (Hoechst, blue) at day 3 post-transfection. Scale, 25 µm. (**M - O**) Quantitative image analysis results. (**P**) Representative immunoblot of mechanical mediators with HSP90 loading control. (**Q**) Protein levels normalized to siControl (number of independent experiments, n = 6 for NUDT5 and pFAK; n = 5 for pMLC2 and ppMLC2). Horizontal lines show medians; error bars show interquartile ranges (**C**) or 95% CI (all others). Statistical significance: Kruskal-Wallis with Dunn’s test (**A**,**B**,**G**,**J**); Mann-Whitney U test (**C**,**D**,**E,H**,**I**,**K**,**M–O**,**Q**).*p < 0.05, **p < 0.01, ***p < 0.001, ****p < 0.0001. n = 3 independent experiments unless noted; color shades denote independent experiments. All data for OVCAR-5-CisR unless indicated.

To corroborate these findings and quantify cellular deformability at the single-cell level, we used quantitative deformability cytometry (q-DC) (*42*); this method uses high-speed imaging of cells as they rapidly deform through the micron-scale pores of a microfluidic device. We previously established that cells that deform more rapidly through the microfluidic pores—or have a lower transit time—tend to have reduced cellular stiffness and increased fluidity (*42*). The q-DC results reveal that siNUDT5 OVCAR-5-CisR cells show a reduction in transit time, as reflected by the downward shift in the distribution of transit times and decreased median transit time for siNUDT5 cells (8.1 ms) compared to siControl (10.0 ms) (p = 1.3 × 10^-8^) (**Fig. 3C**).

Since the mechanical properties of cells are sensitive to their surrounding microenvironment, and in many contexts ovarian tumor cells are adherent to a surface, we measured how NUDT5 impacts the stiffness of cells adhered to a collagen-coated substrate. Force probe indentation of the cytoplasmic region of siNUDT5 OVCAR-5-CisR cells revealed a reduced effective Young’s modulus compared to siControl (median *E_siControl_* = 1100 Pa vs. *E_si-NUDT5_* = 850 Pa, p = 1.2 × 10^-5^) (**Fig. 3D**). The nuclear region also showed a modest but statistically significant reduction in effective Young’s modulus in NUDT5-depleted cells compared to siControl (median *E_siControl_* = 750 Pa vs. *E_siNUDT5_* = 715 Pa, p = 0.025) (**fig. S6A**). Taken together, these results show that depletion of NUDT5 causes cells in both suspended and adherent states to be more deformable, substantiating that NUDT5 contributes to ovarian cancer cell stiffness.

We next asked how the activity of NUDT5 impacts cell mechanical behaviors. ATP is required for actin polymerization and remodeling of the actin cytoskeleton (*43*), so we hypothesized that the enzymatic role of NUDT5 in catalyzing ATP production may contribute to the regulation of cell mechanics. To block NUDT5 activity, we treated cells with the NUDT5 inhibitor (NUDT5-i), TH5427, which binds to NUDT5’s active sites (Trp28 and Trp46) to block ADP-ribose metabolism (*36*). For all subsequent assays, we used inhibitor concentrations from 0.1 to 10 µM for OVCAR-5-CisR cells and 0.01 to 1 µM for PEA-1 cells, which we confirmed do not significantly affect cell viability (OVCAR-5-CisR: p_Veh-10µM_ > 1.0, PEA-1: p_Veh-1µM_ > 1.0) (**fig. S7A-D**). To test whether NUDT5’s effects on cellular mechanics are mediated by its role in ATP production, we first measured intracellular ATP levels following NUDT5 depletion or inhibition. Vehicle-treated OVCAR-5-CisR cells showed ATP levels of ∼12 fmol per cell (**Fig. 3E**), which is consistent with levels reported in MCF7 breast cancer cells (*44*) and within the range reported for mammalian cells more generally (*45*). Pharmacological inhibition of NUDT5 activity reduced cellular ATP levels by 20% (p = 1.5 × 10^-6^ for 1 µM TH5427) and siNUDT5 knockdown by 26% (p = 1.4 × 10^-5^) as measured by a luminescence-based ATP assay (**Fig. 3E**, **fig. S8A**). We confirmed reduced adenosine metabolite levels at the single-cell level using an adenosine-responsive aptamer (*46*), which binds ATP, ADP, AMP, and adenosine (**Fig. 3F-G, fig. S8B-C)**. The reduction in aptamer fluorescence was observed across both whole cell and nuclear regions, consistent with the ATP-specific luminescence measurements. We confirmed using a scrambled non-responsive aptamer as a negative control that the fluorescent signal changes were specific to NUDT5 inhibition and not due to background fluorescence arising from sensor degradation or the delivery or presence of the nonspecific aptamer (**fig. S8D-E**). Consistent with blockade of NUDT5 activity, targeted metabolomics confirmed reduced ATP (p = 0.10) and increased ADP-ribose (p = 0.10), the substrate of NUDT5, with no significant changes in ADP (p = 0.40) or AMP levels (p > 1.0) (**fig. S9A-D**). To test how the ATP-generating activity of NUDT5 compares to the mitochondrial ATP synthase, we treated cells with the ATP synthase inhibitor oligomycin A (*47*). Treatment with oligomycin A similarly caused a reduction in ATP levels (**Fig. 3E**). These findings indicated that NUDT5 contributes to whole cell ATP levels alongside the canonical mitochondrial ATP synthase.

We then tested the effects of NUDT5 blockade on the deformability of OVCAR-5-CisR cells using the filtration assay; these data revealed that NUDT5-i treatment reduced retention, reflecting that NUDT5 activity contributes to regulating the deformability of both OVCAR-5-CisR and PEA-1 ovarian cancer cells (**Fig. 3H-I**). For example, 1 µM of NUDT5-i decreased retention by 14% for OVCAR-5-CisR cells (p = 4.0 × 10^-4^) and by 19% for PEA-1 cells (p = 0.0082). Compared to NUDT5-i, treatment with oligomycin resulted in a greater decrease in retention: 37% reduction for OVCAR-5-CisR cells and 15% for PEA-1 cells. By contrast, treatment with cytochalasin D significantly reduced retention by 95% for OVCAR-5-CisR and by 97% for PEA-1 cells. We confirmed that the altered cell filtration was not due to changes in cell size; cells treated with NUDT5-i have a similar cell size distribution as vehicle-treated cells (**fig. S5C-D**). We corroborated these findings of NUDT5 inhibition by measuring the stiffness of adherent OVCAR-5-CisR cells treated with NUDT5-i (TH5427) using force probe indentation, revealing a 45% reduction in the effective Young’s modulus (*E*) of the cytoplasmic region; cells treated with oligomycin A showed a 49% reduction (median *E_Vehicle_* = 1620 Pa, *E_NUDT5-i_* = 895 Pa, *E_OligA_* = 820 Pa; p_Veh-NUDT5-i_ < 1.0 × 10^-15^, p_Veh-OligA_ < 1.0 × 10^-15^) (**Fig. 3J**). In addition, we observed a similar decrease in the effective Young’s modulus of the nuclear region in OVCAR-5-CisR cells (median *E_Vehicle_* = 910 Pa vs. *E_NUDT5-i_* = 790 Pa, p = 0.0075) (**fig. S6B**). Similar trends were evident with PEA-1 cells (**fig. S6C**). These findings highlight that cellular deformability is regulated by multiple factors: NUDT5 and ATP synthase contribute but F-actin organization has a stronger modulatory role based on the substantially stronger effects of cytochalasin D (**Fig. 3H-I**). We further characterized the effects of NUDT5 inhibition on the cortical tension of OVCAR-5-CisR cells using micropipette aspiration. Consistent with our measurements of cell deformability and stiffness, we found a reduction in cortical tension with NUDT5-i treatment (Vehicle = 0.29 nN/µm vs. NUDT5-i = 0.17 nN/µm, p = 9.0 × 10^-4^) (**Fig. 3K**). Taken together, these findings support that blockade of NUDT5 activity has similar effects as NUDT5 depletion on cell mechanics, with increased activity of NUDT5 causing cells to be stiffer and less deformable.

To gain further insights into how NUDT5 regulates cell mechanical behaviors, we measured the effects of NUDT5 on cell and nuclear shape, which are determined by various factors including structural proteins, intracellular tension, and cell-matrix adhesions. Using confocal microscopy and quantitative image analysis, we found that NUDT5 depletion by siNUDT5 resulted in marked changes in cell morphology, with OVCAR-5-CisR cells showing a more elongated, narrow shape compared to siControl cells (**Fig. 3L, fig. S10**). These changes in cell shape are reflected in the reduction of cell circularity (median circularity of siControl = 0.37 vs. siNUDT5 = 0.21, p < 1.0 × 10^-15^) and cell minor axis length (median length of siControl = 30.19 vs. siNUDT5 = 17.84 µm, p < 1.0 × 10^-15^) (**Fig. 3M**). Analysis of nuclear shape revealed similar differences that tracked with cell shape changes, including a reduction in nuclear circularity with siControl = 0.59 vs. siNUDT5 = 0.55, p < 1.0 × 10^-15^) and nuclear minor axis length (median siControl = 13.73 vs. siNUDT5 = 9.98 µm, p < 1.0 × 10^-15^) (**Fig. 3N**). Similar morphological changes were observed in PEA-1 cells (**fig. S11**). Blockade of NUDT5 had similar effects on cellular and nuclear morphology as NUDT5 knockdown: NUDT5-i treated cells showed an elongated morphology, as reflected by the reductions in cellular circularity and cell minor axis length (**fig. S12**). Inhibition of ATP synthase by oligomycin A also caused cells to become more elongated (**fig. S12**), suggesting this could be a common morphology of starved cells. These global changes in cellular and nuclear shape suggest that NUDT5 contributes to the regulation of both cellular and nuclear morphology.

We next evaluated how NUDT5 impacts the levels and organization of canonical cytoskeletal and nuclear proteins that are determinants of cell and nuclear shape and mechanics. We first assessed F-actin, which is a major determinant of cellular shape and mechanical properties, and found a significant reduction in siNUDT5 cells compared to siControl (median F-actin intensity for siControl = 0.22 vs. siNUDT5 = 0.13, p < 1.0 × 10^-15^) (**Fig. 3O**). Further analysis of other cytoskeletal components showed that siNUDT5 resulted in slightly increased levels of vimentin, as well as non-muscle myosin IIA and IIB (**fig. S13A-C**). By contrast, there was a slight reduction in pMLC and ppMLC2, while no significant difference in the intensity of α-tubulin, suggesting that NUDT5 depletion may alter the levels and organization of intermediate filaments, myosins, and myosin activity, but not levels of microtubules (**fig. S13D-F**). To further define how NUDT5 impacts the activity of key mechanical mediators, we used semi-quantitative immunoblotting. We measured the activity of focal adhesion kinase (FAK), which is central for mechanotransduction and regulation of cell shape, adhesion, and migration (*26*). These data revealed a 20% reduction in phosphorylation of FAK (pFAK) relative to total FAK in siNUDT5 cells (relative expression of pFAK/FAK = 1.00 for si-Control vs. 0.80 for si-NUDT5, p = 0.0022) (**Fig. 3P-Q**). We also evaluated the levels and activity of myosin light chain 2 (MLC2), the phosphorylation status of which reflects the activity of non-muscle myosin II. Depletion of NUDT5 resulted in a marked reduction of pMLC2 and ppMLC2 (**Fig. 3P-Q**), consistent with the immunostaining results (**fig. S13E-F**). Collectively, these findings show that NUDT5 contributes to regulation of the actin cytoskeleton and focal adhesions, which are central in mediating cell mechanical behaviors such as cell shape, stiffness, cell-matrix interactions, and force generation.

### Functional relevance of NUDT5 in ovarian cancer progression

To explore the relevance of NUDT5 in ovarian cancer, we analyzed NUDT5 expression across tumor and normal tissues by mining data from The Cancer Genome Atlas (TCGA). NUDT5 levels were elevated in multiple tumor types; ovarian tumors showed a ∼2.7-fold increase in NUDT5 levels compared to normal ovarian epithelium (**Fig. 4A**). Using *in situ* hybridization of a patient tissue microarray (N = 42), we found that NUDT5 transcripts were localized to the epithelial region of HGSOC tumors (**Fig. 4B-C**). Importantly, analysis of gene expression data of epithelial ovarian cancers (*48*) reveals that higher NUDT5 levels were associated with higher tumor grade (**Fig. 4D**) as well as reduced overall and progression-free survival (**Fig. 4E, fig. S14A**).

**Fig. 4.**
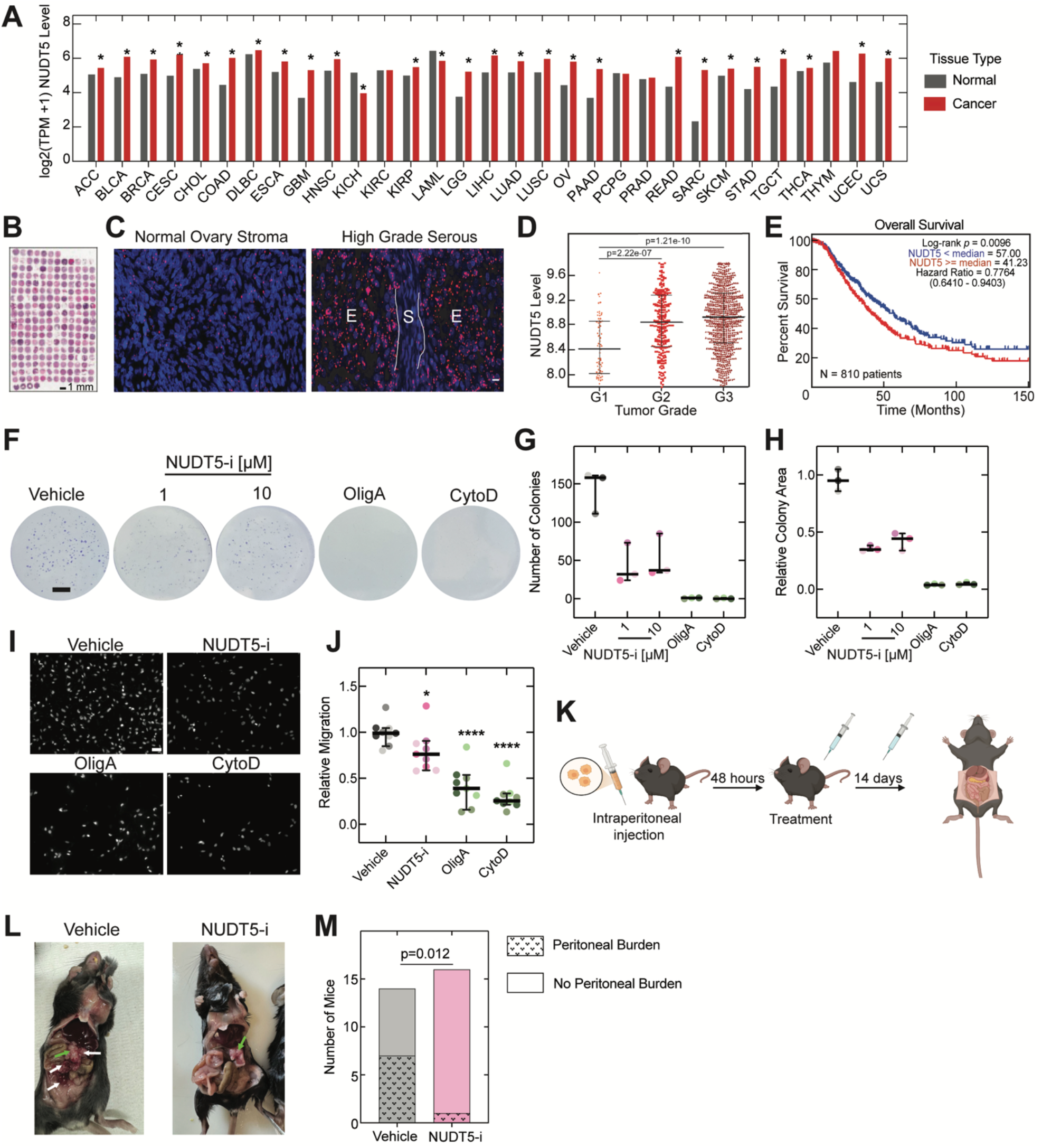
Functional relevance of NUDT5 in ovarian cancer progression. (**A**) Median log2(transcripts per million +1) NUDT5 levels across TCGA tumors versus GTEx normal tissues using DESeq2; ovarian cancer, OV. * p_adj_< 0.05, Benjamini and Hochberg false discovery rate. Data from the Cancer Genome Atlas (TCGA) and Genotype-Tissue Expression (GTEx) Portal. (**B**) H&E staining of tissue microarray of benign and high-grade serous carcinoma tissues (N = 42 patients). (**C**) Representative images from the RNAScope *in situ* hybridization of NUDT5 on the tissue microarray showing normal ovarian stroma and high-grade serous carcinoma with stromal (S) and epithelial (E) cells. Scale, 25 µm. (**D, E**) Association of NUDT5 levels in epithelial ovarian carcinomas with: tumor grade (**D**) and overall patient survival (**E**). Data obtained and visualized using CSIOVDB (*48*). (**F**) Representative images of OVCAR-5-CisR colonies formed on tissue culture plates at 10 d after daily treatment with vehicle (DMSO), NUDT5-i (TH5427), oligomycin A (OligA), or cytochalasin D (CytoD). Scale, 1 cm. (**G, H**) Quantification of colonies. Horizontal lines show median and error bars represent 95% CI (n = 3 independent experiments). (**I**) Representative images of DNA-labeled (Hoechst 33342) OVCAR-5-CisR cells that migrated through 5 µm pores in a transwell migration assay after 24 h (24 h treatment with vehicle, 10 µM NUDT5-i, 1 µM OligA, or 75 nM CytoD. Scale, 50 μm. (**J**) Quantification of the number of migrated cells. Horizontal lines show median ± 95% CI.; each dot represents an individual image from one of n = 3 independent experiments, which are represented by different colors. Mann-Whitney U test, *p < 0.05. (**K**) Workflow schematic to evaluate NUDT5-i effects on ovarian cancer cell dissemination *in vivo*. Created with Biorender.com. (**L**) Representative images of mice after 14 d treatment with vehicle (sterile water) or NUDT5-i; green arrows denote nodules in the omentum, white arrows denote nodules in the peritoneal cavity. (**M**) Quantification of peritoneal burden (n = 3 independent experiments); Chi-Square (χ^2^) test, p = 0.012.

We next sought to assess the functional role of NUDT5 in cancer progression using *in vitro* colony formation and migration assays. The ability of cells to form colonies *in vitro* is a hallmark of cancer cells that is associated with the propensity of cells to form tumors *in vivo*. Colony formation also depends on mechanical behaviors of cells, such as motility and adhesion (*49*). Quantification of the number and size of colonies at ten days after cell seeding revealed that NUDT5-i treatment resulted in a marked reduction in the total area of colonies (90% reduction with 1 µM TH5427, and 84% reduction with 10 µM TH5427; p = 0.0017) (**Fig. 4F-H**). By comparison, blockade of ATP synthase activity with oligomycin A or perturbation of F-actin with cytochalasin D resulted in negligible colony formation, reflecting the importance of ATP and actin remodeling for cancer cells to form colonies.

Cell migration is another *in vitro* hallmark of cancer that is indicative of metastatic potential. To evaluate the effects of NUDT5 in mediating the confined migration of ovarian cancer cells, we performed a transwell assay using a membrane with 8 µm pores. NUDT5-i treatment significantly reduced transwell migration by 23% (p = 0.01) compared to the vehicle control, showing that NUDT5 activity contributes to the migratory capacity of ovarian cancer cells (**Fig. 4I-J**). Similar to the colony formation results, while NUDT5 contributes to regulating cell migration, the blockade of ATP synthase and F-actin remodeling had stronger effects, consistent with the more modest contributions of NUDT5 to modulating cellular ATP and cytoskeletal actin.

To test whether NUDT5 inhibition affects ovarian cancer progression *in vivo*, we transplanted SO ovarian cancer cells into syngeneic (C57BL/6) mice via intraperitoneal injection, treated the mice with either NUDT5-i or vehicle control, and assessed tumor involvement of the omentum, the preferred site of ovarian cancer metastasis (*50*), as well as peritoneal burden at sites outside than the omentum after 14 days (**Fig. 4K**). The SO cell line exhibits highly aggressive behavior, characterized by rapid, extensive intraperitoneal tumor spread (*51*). While mice treated with NUDT5-i showed some tumor formation in the omentum (**fig. S14B**), we found that NUDT5-i treatment significantly reduced peritoneal tumor burden, with only one out of 16 mice showing extensive tumor burden beyond the omentum compared with seven out of 14 vehicle-treated control mice (**Fig. 4L-M**). These findings suggest that NUDT5 inhibition can reduce ovarian cancer dissemination *in vivo* and support further investigation of NUDT5 as a therapeutic target.

## DISCUSSION

Here, we established deformability-based screening as a platform for the systematic discovery of molecular mediators of cellular mechanical behaviors. Screening 1,280 bioactive compounds identified 92 modulators of cellular deformability. Transcriptomic connectivity analysis of 21 top compounds revealed NUDT5, a Nudix hydrolase, as a previously unrecognized mechanical mediator that controls cellular stiffness, intracellular tension, morphology, and migration through ATP production. In ovarian cancer models, NUDT5 inhibition reduced colony formation, transwell migration, and peritoneal tumor burden, thereby establishing proof-of-concept for the deformability screening pipeline as a tool for unbiased discovery of mechanical mediators that may have translational potential.

Our findings validate cellular filtration as a screening platform to identify mechanical mediators or ‘mechanomodulators’. The screen preferentially identified compounds that are cytoskeletal and ECM-targeting agents (∼9-fold enrichment compared to original LOPAC library).

All top six hits reduced cellular deformability as well as migration and invasion, including the microtubule-targeting agents vincristine and vinblastine (*20, 21*), along with wiskostatin, which inhibits the Arp2/3 complex, which is well known to modulate actin architecture, cell motility, and cell stiffness (*18, 52*). Further transcriptomic connectivity analysis across 21 top compounds (Z > 3) revealed known mediators of cell mechanics (KRAS, PTK2, EGFR) (*7, 25–27*) in addition to NUDT5, which we define here as a mechanical mediator. Interestingly, the top 92 compounds that significantly reduced cell filtration also include drugs that are implicated in cell stress and neurotransmission. Further study of these compounds could provide additional insights into the intersection of mechanobiology with other established signaling pathways.

The deformability screening platform offers distinct advantages over existing strategies. Unlike AFM-based screens, which require specialized equipment and sequential single-cell measurements, the cellular filtration assay is compatible with standard plate readers and could be further scaled for even larger screens. A key innovation enabled by the scale of the deformability screen is the generation of datasets of mechanomodulating compounds that enable computational prediction of mechanical mediators. From the 92 hits identified, we prioritized 21 compounds (Z > 3) for transcriptomic connectivity analysis using the Connectivity Map (cMAP) database to predict potential mechanical mediators, which revealed NUDT5. By identifying consensus transcriptional signatures across multiple compounds, this approach could narrow the search space for mechanical mediators compared to genome-wide genetic screens. While lower-throughput methods such as AFM are valuable for mechanistic studies, cell filtration enabled us to simultaneously generate datasets across dozens of compounds on practical timescales, which enabled robust computational predictions. One caveat is that deformability-based screening measures bulk populations of cells; single-cell resolution will be important to resolve cell-to-cell variability and subpopulations of cells with distinct mechanical phenotypes. We envision that deformability screening can complement other strategies to identify mechanical mediators at the single-cell level such as mechanical phenotyping coupled to transcriptomics (*53*) and *de novo* network-based inference (*54*). Together, integrated screening of cells based on mechanical phenotypes coupled with omics-level analyses can provide a scalable framework for systematic mapping of the ‘mechanome’—the set of genes and pathways that regulate cellular mechanical behaviors.

In this study, we define the role of NUDT5 in cellular mechanobiology. NUDT5 contributes to cellular mechanical properties—including deformability, stiffness, and cortical tension—as well as morphology and confined migration. Mechanistically, NUDT5 depletion reduced levels of F-actin (by 41%), pMLC2/MLC2 (by 39%), ppMLC2/MLC2 (by 46%), as well as pFAK/FAK (by 20%), consistent with reduced cell stiffness and intracellular tension. While we found that NUDT5 depletion resulted in a modest increase at the protein level of vimentin and non-muscle myosin IIA and IIB, this could reflect a compensatory response to the loss of F-actin. Depletion of NUDT5 also resulted in transcription-level changes that could contribute to the observed altered mechanical behaviors. For example, NUDT5 depletion resulted in differential expression of RhoGTPases and integrins, which may also contribute to the observed changes in F-actin, cellular tension, and mechanical properties including cell shape and stiffness.

We hypothesized that NUDT5 contributes to mediating cell mechanical behaviors via its role in regulating cellular ATP levels, whereby NUDT5-catalyzed ATP production directly supports cytoskeletal remodeling and cellular force generation. Depletion or blockade of NUDT5 reduced cellular and nuclear ATP levels by ∼20%, although to a lesser degree than the ∼44% reduction in ATP caused by inhibition of mitochondrial ATP synthase. Despite producing a smaller reduction in ATP levels (∼20% vs ∼44% with ATP synthase inhibition), blockade of NUDT5 resulted in similar changes in cell morphology and deformability, supporting the importance of NUDT5-regulated ATP in modulating cell mechanical behaviors. Notably, while partial ATP depletion softened cells in our experiments, complete ATP depletion causes cell stiffening or ‘rigor mortis’ (*55*); this highlights the complex, non-linear relationship between ATP and cell mechanical behaviors.

Since actin polymerization consumes ATP, the reduction in ATP availability with NUDT5 depletion or inhibition could directly impair actin remodeling; this is consistent with our results showing reduced filamentous actin and increased cell deformability. Although the proportion of total cellular ATP consumed by cytoskeletal remodeling remains unresolved and appears to depend on the measurement approach used (*56–58*), our data indicate that the ∼20% reduction in ATP following NUDT5 depletion or inhibition is associated with altered actin organization and reduced cell stiffness. However, because NUDT5 depletion also elicits broad transcriptional changes, we cannot exclude that these gene expression changes contribute to the observed mechanical phenotype alongside, or independently of, reduced ATP availability.

Beyond ATP-depletion effects, changes in adenosine nucleotide ratios can also alter cytoskeletal dynamics. For example, an increased AMP:ATP ratio due to reduction in ATP could trigger AMP-activated protein kinase (AMPK) activation, which can have downstream effects via RhoA/ROCK to modulate cell mechanical behaviors via MLC2 phosphorylation and F-actin remodeling (*59*); however, rather than the increased MLC2 phosphorylation that AMPK/RhoA/ROCK activation would predict, we found that NUDT5 depletion reduced pMLC2, ppMLC2, and F-actin, and cell stiffness, suggesting that the direct effects of ATP depletion dominate over AMPK-mediated signaling in NUDT5 regulation of cell mechanics. Beyond nucleotide-mediated effects on cell mechanics, chromatin organization can also mediate cell mechanical behaviors (*60*), and prior work shows that NUDT5 regulates chromatin remodeling (*31*); how NUDT5 modulates chromatin architecture in ovarian cancer cells requires further investigation.

Taken together, we hypothesize that NUDT5 functions as a mediator of cell mechanical behaviors through complementary mechanisms that span timescales: on short timescales of seconds to minutes, NUDT5-mediated ATP could enable actomyosin remodeling, modulation of cellular force generation, and chromatin remodeling to support transcriptional reprogramming; NUDT5-dependent changes in ADP-ribose levels may also independently contribute to chromatin remodeling on these short timescales (*61*). On longer timescales of hours, changes in NUDT5 levels could drive transcriptional changes that regulate cell mechanical behaviors, for example, through altered expression of Rho GTPases and integrins. Dissecting the relative contributions of these parallel pathways, including acute ATP-dependent effects, ADP-ribose dependent effects, and longer-term remodeling resulting from transcriptional changes, will require temporal studies with inducible NUDT5 perturbation. While our findings substantiate that the enzymatic activity of NUDT5 contributes to cellular mechanical behaviors, it will be important in future work to investigate the non-enzymatic role of NUDT5 in regulating nucleotide homeostasis (*33, 34*), as well as to disentangle ATP-dependent versus ADP-ribose-dependent contributions to NUDT5’s effects on cell mechanics. Studies of catalytically inactive NUDT5 mutants would further delineate the enzymatic versus non-enzymatic contributions of NUDT5 as a mechanical mediator.

The role of NUDT5 in regulating cellular ATP production could have functional consequences for nuclear mechanics and cell migration. Nuclear-localized NUDT5 could provide a local source of ATP to support nuclear remodeling; since the nucleus rate-limits the confined migration of cells (*62, 63*), we speculate that NUDT5 enables the confined migration of cells by supplying the ATP required for cells to rapidly remodel their nuclei to deform through the confined spaces of dense extracellular matrix (*64*). NUDT5 may also contribute to ATP-mediated DNA repair of double-strand DNA breaks that can accumulate during confined migration (*65*), thereby maintaining genomic integrity. An important unresolved question is whether mechanical stress directly modulates NUDT5 levels, activity, or localization, which could complement AMPK in supporting cellular adaptation to fluid shear stresses (*66*) or phospho-creatine (pCr)/creatine kinase (CK) in supporting ATP recycling for cells on increased matrix stiffness (*67*). Future studies will disentangle the mechanisms through which NUDT5 could provide a selective advantage for cells to respond and adapt to mechanical stresses.

Our findings also point to the potential role of NUDT5 in ovarian cancer. Higher NUDT5 levels are associated with increased tumor grade and worse patient survival, suggesting the potential of NUDT5 as a biomarker for ovarian cancer progression. *In vitro*, NUDT5 modulates cell behaviors that are relevant for cancer, such as deformability, confined migration, and colony formation. The initial *in vivo* studies show that NUDT5 inhibition reduces peritoneal spread, suggesting that NUDT5 blockade reduces cell-cell and cell-matrix interactions that are important for colony formation and peritoneal colonization. While our *in vivo* findings establish proof-of-concept for NUDT5 as a therapeutic target, the mechanisms by which NUDT5 inhibition reduces peritoneal spread remain to be fully elucidated. Further evaluation of NUDT5 inhibition in combination with standard-of-care therapies and in additional ovarian cancer models, such as patient-derived xenografts, is also needed. As a regulator of cellular mechanobiology, ATP production, and DNA repair, NUDT5 sits at the intersection of multiple processes that are crucial for chemoresistance and disease spread, and may therefore be positioned as a promising therapeutic target for platinum-resistant ovarian cancers.

Beyond NUDT5, the deformability screening platform identified compounds that suppress cancer cell invasion and migration without compromising cell viability, highlighting the potential of ‘mechanomodulatory’ therapeutics to complement—and even synergize with—conventional cytotoxic therapies while potentially reducing side effects. Among the top six hits, four compounds reduced migration and invasion while maintaining > 90% cell viability at effective concentrations; NUDT5 inhibition similarly reduced migration without cytotoxic effects. By targeting mechanical behaviors rather than cell viability or specific mutations, mechanomodulators offer a complementary strategy to existing cancer therapeutics. The potential of mechanomodulatory compounds for blocking cancer progression is supported by previous reports of the small molecule 4-HAP, which increases cancer cell stiffness, reduces cell motility, and blocks metastasis in mouse models of both pancreatic and colon cancer (*68–70*). More recently, the FDA approval of FAK inhibitors for treating low-grade serous ovarian cancer demonstrates the clinical viability of targeting mechanomodulatory pathways. Our finding that NUDT5 depletion reduced pFAK levels by 20%, alongside reductions in pMLC2 and ppMLC2, suggests that NUDT5 inhibition could complement FAK-targeted therapies (*71*).

Screening for drugs based on deformability may also circumvent the genomic heterogeneity that limits mutation-targeted approaches. Cell mechanical phenotype represents a convergent property across genetically diverse tumors: drug-resistant, mesenchymal-type ovarian cancer cells are consistently more deformable than their drug-sensitive, epithelial-type counterparts regardless of their mutational landscape (*7*). While the relationship between mechanical phenotype and invasive behavior can vary by cancer type and microenvironmental context, altered deformability is associated with invasive and metastatic phenotypes across multiple malignancies (*6, 13, 18*), suggesting that mechanomodulation may have broad applicability. Mechanomodulators may thus represent a conceptually distinct therapeutic paradigm, modulating cell mechanical behaviors towards less invasive phenotypes rather than killing cells or targeting specific mutations. This approach could be particularly valuable as a maintenance strategy to prevent metastatic dissemination in patients with chemotherapy-resistant cancers or minimal residual disease, where residual tumor cells have already acquired resistance to conventional cytotoxic and targeted therapies.

Together, this work establishes cellular deformability screening as a platform for the discovery of mechanical mediators and validates mechanomodulation as a potential therapeutic strategy. More broadly, systematic mapping of the mechanome addresses a fundamental gap in cell biology: while comprehensive maps of the genome, transcriptome, and proteome provide reference frameworks for understanding cellular function, equivalent systems-level knowledge of the genes controlling mechanical phenotypes remains incomplete. Cellular mechanical behaviors—stiffness, deformability, shape control—are increasingly recognized as disease biomarkers and therapeutic targets across cancer, fibrosis, immune dysfunction, and developmental disorders. Given that cell filtration can be applied across cell types from fibroblasts to immune cells (*19, 72*), the deformability screening platform could have applications beyond cancer for diseases with mechanobiological dysfunction. By systematically mapping molecular mediators to cellular mechanical phenotypes, this approach has potential to advance both fundamental mechanobiology and deliver new strategies for translational ‘mechano-medicine’ across diseases where mechanical dysfunction drives pathology.

## Materials and Methods Experimental Design

### Cell culture

Human cisplatin-resistant ovarian cancer (OVCAR-5-CisR) cells were cultured in DMEM (+L-Glutamine, +Glucose, +Sodium Pyruvate, Gibco) supplemented with 10% fetal bovine serum (FBS, Gemini), 1% penicillin-streptomycin (VWR), and 10 µM cisplatin (Sigma-Aldrich) as previously described (*73*). PEA-1 cells were cultured in DMEM media with 10% FBS and 1% penicillin-streptomycin. Cells were maintained at 37 °C with 5% CO_2_ and tested for mycoplasma routinely. Cell identity was confirmed by short tandem repeat (STR) profiling.

### Drug treatments

The library of pharmacologically active compounds (LOPAC) (LO1280) was purchased from Sigma. Vinblastine (V1377), vincristine (V8879), SCH 58261 (S4568), bezafibrate (B7273), ganciclovir (PHR1593), wiskostatin (W2270), paclitaxel (T7402), cytochalasin-D (C8273), and oligomycin A (75351) were from Sigma-Aldrich. NUDT5 inhibitor (NUDT5-i, TH5427) was obtained from Tocris (6534). Drugs were reconstituted in dimethyl sulfoxide (DMSO) to generate 10 mM stock solutions according to manufacturers’ instructions. All drug treatments were prepared in DMSO at a final concentration of ≤ 0.1%; we used 0.1% DMSO as the vehicle control for all *in vitro* experiments. For *in vivo* experiments, NUDT5-i was reconstituted in sterile water and sterile water served as the vehicle control.

### Cell filtration assay

To screen cells based on cellular deformability, we used cell filtration as previously described (*7*) and optimized the assay to have *Z’* ≥ 0.5, where *Z’ = 1 – (3σ_Positive_+3σ_Negative_) / |µ_Positive_-µ_Negative_|.* OVCAR-5-CisR cells were treated with the LOPAC compounds at a concentration of 10 µM for 24 h. Prior to the deformability assay, cells were washed with 1× Phosphate-Buffered Saline (PBS, DNase-, RNase-& Protease-free, Mediatech), treated with trypsin, and resuspended in fresh medium to a density of 5 × 10^5^ cells/mL. We loaded 350 µL of cell suspension into the loading chamber of the filtration device with a polycarbonate membrane with 10 µm pore size (Millipore) and applied 1.4 kPa of air pressure for 15 sec using a pressurized air tank. To quantify the retained volume after filtration, we transferred the cell suspension remaining in the top well to a 96-well plate and measured the absorbance of phenol red at 560 nm using a plate reader (Infinite M1000, Tecan). To calculate the Z-score, we normalized absorbance relative to the vehicle (DMSO) control across each plate.

We also used cell filtration to validate NUDT5 as a mechanical mediator. PEA-1 and OVCAR-5-CisR cells were treated with siNUDT5 or TH5427 for 24 h. Prior to filtration, we measured the number and size distribution of cells using an automated cell counter (TC20, BioRad). We loaded 400 µL of cell suspensions at a density of 5 × 10^5^ cells/mL into the loading chamber, applied air pressure of 2.3 kPa for 30 sec (OVCAR-5-CisR) or 5.5 kPa for 60 sec (PEA-1), and measured the absorbance of the retained cell suspension (560 nm) using a plate reader (SpectraMax M2, Molecular Devices) to quantify the retention volume.

### Drug enrichment analysis

Broader functional categories were generated from drug class annotations provided by the manufacturer (Sigma) (**fig. S1A**). For each functional category, we determined the log2FC based on the ratio of compounds from the 92 hits relative to the 1280 drug library. Enrichment analysis was conducted using the drugSEA function from the DMEA R package, implementing a rank-based enrichment framework adapted for drug class annotations (*74*). Statistical significance was determined by permutation testing to calculate the p-adjusted value. Analysis was performed using in R version 4.5.2.

### Cell viability assays

To evaluate the top six hits for cytotoxicity, we plated cells in a 96-well plate (3,000 cells per well), treated them with 1 nM to 10 µM of drug or vehicle (DMSO) control, and incubated at 37 °C, 5% CO_2_. After 48 h, CellTiter-Glo Reagent (Promega) was used to determine cell viability according to the manufacturer’s instructions (GloMax 20/20 Luminometer, Promega). A higher luminescence indicates a larger number of viable cells. Percent viability was determined by normalizing to vehicle (DMSO)-treated cells.

To determine the effects of the NUDT5 inhibitor, TH5427, on the viability of OVCAR-5-CisR and PEA-1 cells, we used MTT and trypan blue assays. For the MTT assay (Abcam), we treated 7,500 cells per well of a 96-well plate with 1 nM to 100 µM TH5427 for 24 hours. The drug was then removed, replaced with 100 µL serum-free media and 100 µL MTT reagent, and incubated for 3 hours at 37°C. After incubation, the media and reagents were removed and 150 µL of MTT solvent was added for 15 minutes on an orbital shaker. Absorbance was read at 590 nm using a plate reader (SpectraMax M2, Molecular Devices). Because MTT assays are dependent on intracellular ATP levels, which NUDT5 modulates, we confirmed cell viability using trypan blue exclusion. To assess viability using trypan blue, we plated 100,000 cells per well of a 6-well plate. Following attachment, cells were treated with NUDT5 inhibitor at various concentrations for 24 – 72 h. Following treatment and detachment, cells were stained with 0.4% trypan blue (BioRad). Viable cells were counted using a TC20 (BioRad).

### Cell cycle assay

To perform cell cycle analysis, we treated 0.3 × 10^6^ cells per well of a 6-well plate with drugs for 24 hours prior to harvesting and resuspending in fresh medium to a density of 2 × 10^6^ cells/mL. Cells were washed once in PBS containing 1% FBS (Gibco) by centrifugation and resuspended in 70% ethanol (Fisher Scientific) solution made in PBS. Cells were fixed in the ethanol solution overnight at -20 °C, and then washed once in PBS prior to staining with propidium iodide (PI) (50 µg/mL, Thermo Fisher Scientific) and 2.5 mg/mL RNase solution (Invitrogen) in PBS) for 30 min at 37 °C. To minimize cell aggregation, cell suspensions were passed through a cell strainer with 35 µm mesh size (BD Falcon) prior to analysis using flow cytometry (LSRFortessa cell analyzer, BD Falcon).

### *In vitro* invasion & migration assays

To test effects of the top six compounds on cell invasion and migration, we used a transwell assay equipped with polycarbonate membranes with 8 µm diameter pores (Culturex, R&D Systems). Invasion was measured using the same assay with basement membrane extract (BME) (Culturex, R&D Systems). Cells were pre-treated with drugs for 24 h. We plated 3,000 cells per well in each transwell chamber of a 96-well plate and added cell culture media containing the indicated concentration of drug; the same drug-containing media was also added to the bottom wells. After incubation for 24 h at 37°C and 5% CO_2_, cells were stained with Calcein AM, and the number of cells was quantified using a plate reader to measure fluorescence (485 nm excitation, 520 nm emission) (Infinite M1000, Tecan). The percentage of migrated/invaded cells was calculated by normalizing to vehicle (DMSO)-treated cells.

To measure the effects of NUDT5 on cell migration, we used a transwell assay that contained membranes with 5 µm diameter pores (Corning, 3421). We seeded 50,000 cells per transwell chamber inserts of a 24-well plate in culture media. After 24 h for cell attachment, cells were serum starved and culture medium containing 20% FBS was placed in the basal chamber. After 24 h incubation at 37°C and 5% CO_2_, the non-migrated cells were removed from the top-side of the membrane facing the apical chamber. Then, cells were washed with 1x PBS followed by fixation with 4% paraformaldehyde. DNA was visualized using Hoechst (1:1000) and imaged using an inverted fluorescence microscope (Zeiss Axio Observer) equipped with a 10× objective (N.A. 0.3). We quantified the number of nuclei using Fiji (ImageJ) to measure the number of migrated cells.

### Transcriptomic connectivity analysis

Transcriptomic connectivity between drug treatment signatures and gene perturbation signatures was analyzed using the Touchstone tool in CLUE (CMap Linked User Environment; https://clue.io/touchstone; inquiry date 1/6/2026), a cloud-based platform for querying perturbational data from the Library of Integrated Network-based Cellular Signatures (LINCS) project (*23*).

### Small interfering RNA (siRNA) knockdown

For siNUDT5 experiments, OVCAR-5-CisR and PEA-1 cells were plated in a 6-well dish at a density of 300,000 cells per well. The following day, RNA interference was performed using a Non-targeting Pool siRNA (Qiagen, 1027281) or NUDT5 siRNA (Qiagen, 1027418, GeneGlobe ID: SI04300128|S2; denoted as siNUDT5 #1 or Integrated DNA Technologies, as previously described (*36*); denoted as siNUDT5 #2), and transfections were carried out using Lipofectamine 3000 (Invitrogen, L3000001) according to the manufacturer’s protocol. Briefly, 7.5 µL of Lipofectamine was mixed with 125 µL of Opti-MEM (ThermoFisher, 31985062). Concurrently, 7.5 µL of 10 µM siRNA was mixed with 125 µL of Opti-MEM. These two solutions were mixed and incubated at room temperature for 15 min. The siRNA-lipid complexes were then added dropwise to cells in 3 mL of media and incubated under culture conditions. Samples were collected at 48-72 hours after transfection for qPCR and protein expression analysis to confirm knockdown. Corresponding siRNA sequences can be found in **Table S2**.

### RNA sequencing and analysis

RNA was extracted using TRIzol reagent (Thermo Fisher Scientific, 15596026) and the RNeasy Mini Kit (Qiagen, 74104) according to the manufacturer’s instructions. RNA quality and concentration were assessed by spectrophotometry (NanoDrop 8000, Thermo Fisher Scientific). Bulk RNA sequencing was performed on OVCAR-5-CisR cells 48 h after siRNA-mediated NUDT5 knockdown. Library preparation and sequencing were performed at the UCLA Technology Center for Genomics and Bioinformatics (TCGB). Libraries for RNA-Seq were prepared with KAPA Stranded RNA-Seq Kit with RiboErase Kit. Ribosomal RNA was depleted by hybridization of complementary DNA oligonucleotides, followed by treatment with RNase H and DNase and RNA fragmentation. First-strand cDNA synthesis was performed using random priming, followed by second-strand synthesis to convert cDNA:RNA hybrid to double-stranded cDNA and incorporates dUTP into the second cDNA strand. cDNA generation was followed by A-tailing, adaptor ligation, and PCR amplification. Different adapters were used for multiplexing samples in one lane. Sequencing was performed on Illumina NovaSeq X Plus for PE 2×50 bp run. Data quality was assessed using the Illumina Sequence Analysis Viewer. Demultiplexing was performed with Illumina Bcl2fastq software (v2.19.1.403). Alignment was performed using STAR (*75*) with human reference genome GRCh38. The Ensembl Transcripts release GRCh38.108 GTF was used for gene feature annotation. For normalization of transcripts counts, counts per million (CPM) normalized counts were generated by adding 1.0E-4 followed by CPM normalization. Sequencing depth averaged 1250M/lane. Differential gene expression analysis was conducted using DESeq2 (version 1.48.2). Genes with Benjamini-Hochberg adjusted p-value, p_adj_ < 0.05, were considered differentially expressed. Overrepresentation analysis (ORA) and gene set enrichment analysis (GSEA) were performed using clusterProfiler (version 4.16.0) against Gene Ontology (GO) Biological Processes gene sets with org.Hs.eg.db used for annotation. Biological processes with p_adj_ < 0.05 were considered significant. For GSEA, genes were ranked based on log_2_ fold change.

### Quantitative reverse transcriptase-polymerase chain reaction (qRT-PCR)

To quantify mRNA levels of NUDT5, we used qRT-PCR. In brief, we extracted RNA from cells using Trizol® (Invitrogen, 15596026) and the PureLink RNA mini kit (Invitrogen, 12183025) according to the manufacturer’s instructions. For cDNA synthesis, 500 ng of RNA was reverse transcribed using the Maxima First Strand cDNA Synthesis Kit (ThermoFisher Scientific). Quantitative PCR was performed using Maxima SYBR Green/Fluorescein qPCR Master Mix (ThermoFisher Scientific) on a CFX qPCR system (Bio-Rad). qRT-PCR data were analyzed with CFX Manager 3.1 (Bio-Rad) and gene expression levels were normalized to *PUM1*. Primers used for qRT-PCR are included in **Table S3**.

### Quantitative deformability cytometry (q-DC)

To measure cellular deformability at the single-cell level, we used q-DC as previously described (*42*). In brief, microfluidic devices were fabricated using standard soft lithography methods. Polydimethylsiloxane (PDMS) was poured at a 10:1 w/w base-to-crosslinker ratio, cured, and bonded to a No. 1.5 glass coverslip (Thermo Fisher Scientific) using plasma treatment (Plasma Etch). Within 48 hours of device fabrication, suspensions of OVCAR-5-CisR cells at a concentration of 2 × 10^6^ cells/mL were driven to deform through constrictions of 9 μm (width) × 10 μm (height) by applying 69 kPa of air pressure across the device. We captured images of cell shape during flow-induced cell deformation on the millisecond timescale using a CMOS camera with a capture rate of 1600 frames/s (Vision Research) mounted on an inverted Axiovert microscope (Zeiss). Using a MATLAB (MathWorks) code (https://github.com/rowatlab), we analyzed the time-dependent shape and position changes of individual cells as they deform through a micron-scale gap in the microfluidic device; transit time was measured for individual cells as a function of cell diameter as previously described (*42*); in brief, this is the timescale for the leading edge (front) of a cell to travel through the constriction from the base line to exit line. Single-cell data was size-gated at +/-3 µm or +/-4 µm relative to the median pooled cell size of 24 µm.

### Force probe indentation

To measure cellular stiffness, we used force probe indentation (Pavone, Optics11 Life) equipped with a probe (3 µm radius, 0.02 N/m stiffness). The probe was calibrated in cell culture medium, and all force probe measurements were conducted on cells submerged in cell culture medium with 25 mM HEPES buffer. Cells were treated with TH5427 (1 - 10 µM for OVCAR-5 cells or 0.1 - 1 µM for PEA-1 cells), oligomycin A (1 µM) and cytochalasin D (75 nM) for 24 h before measurements. We used phase contrast microscopy to ensure that force probing was applied to either the cytoplasmic or nuclear region. We measured the displacement to quantify the effective Young’s modulus by fitting the Hertzian model to the force-indentation curves up to 16% of the probe tip radius, or 0.48 µm; this is the effective length scale over which we measure cell stiffness. The Hertzian model considers the probe as a parabolic indenter, which deforms the cell (*76*) with a probe indentation depth ℎ and applied force *F = 4/3(E)/(1-v*^2^*) √R* ℎ^3/2^ where *E* is the Young’s modulus of the sample, *R* is the radius of curvature of the probe tip, and *v* is the sample’s Poisson’s ratio, which is typically assumed to be 0 ≤ *v* ≤ 0.5 for biological materials (*76, 77*). For small ℎ/*R* ratios (e.g., 0.16) and *E_effective_* >> *E_cell_*, the sample Young’s modulus *E* is related to the effective Young’s modulus *E_effective_* as *E = E_effective_(*1-v^2^) (*77*).

### Micropipette aspiration

To measure cortical tension, we use a micropipette aspiration setup as previously described (*78*). Micropipettes of ∼7.5 μm diameter (exact diameter was measured for each pipette during use) were stabilized at the bottom of the cell chamber. We seeded 300,000 cells/well in a 6 well tissue culture treated plate and allowed them to adhere for 24 h before treatment with 1 µM TH5427 for 24 hrs. Prior to measurement, cells were detached with trypsin, suspended in culture medium, and placed in the imaging chamber. A small negative aspiration pressure was generated to secure the cell in the pipette tip without causing deformation of the cell cortex. The cell was then raised off the surface, and the aspiration pressure was increased to the equilibrium pressure (*ΔP*) at which the length of the cell inside the pipette (*L_p_*) is equal to the radius of the pipette (*R_p_*). Effective cortical tension (1 nN/µm = 0.001 Pa) was quantified using the Young-Laplace equation: *ΔP = 2T_eff_ (1/R_p_ - 1/R_c_)*, where *ΔP* = aspiration pressure that produced a deformation where *L_p_ = R_p_*, *T_eff_* = effective cortical tension, *R_p_* = radius of the pipette, and *R_c_* = radius of the cell outside the pipette.

### Immunostaining and imaging

For immunostaining, cells were fixed with 4% paraformaldehyde (Electron Microscopy Sciences) for 10 minutes, washed twice with PBS and permeabilized with 0.5% Triton-X-100 (Sigma), followed by two washes with PBS. Samples were blocked with 5% donkey serum (Jackson Immunoresearch) in PBS for 30 minutes and incubated with primary antibodies (as detailed in **Table S4**) overnight at 4°C. The next day, the samples were washed twice with PBS with 0.05% Tween-20 and once with PBS, followed by a 1-hour incubation with Alexa 488 and/or Alexa 546-labeled secondary antibodies (Molecular Probes) and fluorescein isothiocyanate-conjugated phalloidin (Invitrogen, R37112) and 4,6-diamino-2-phenylindole (DAPI) (Invitrogen, D3571) to label the actin-cytoskeleton and nuclei, respectively. Epifluorescence images were collected using a Zeiss Axio Observer Z1 with a 20× objective (N.A. 0.8) or 40× objective (N.A. 0.95); confocal images were acquired using a Leica SP8 Confocal Laser Scanning microscope equipped with a 40× objective (N.A. 1.25).

### Quantitative image analysis

For image analysis, cell and nuclear masks were first created using images of phalloidin and DAPI-labelled cells as templates (CellPose). Cell and nuclear morphology and fluorescence intensity measurements were quantified using a CellProfiler (v. 4.2.1) macro, where the masks were applied to the corresponding stain image to measure the size and shape of each cell and nucleus as well as the average and integrated fluorescence intensity within each cell and nucleus.

We used a similar approach to quantify fluorescence levels of the adenosine-responsive aptamer, where cellular adenosine metabolites refer to the fluorescence across the whole cell and nuclear adenosine metabolites is defined as fluorescence in the nuclear region. The derived integrated fluorescence intensity values for both the whole cell and nuclear regions were normalized to the median fluorescence intensity of a scrambled non-responsive aptamer for each corresponding condition (**fig. S8D-E**) to obtain the final fluorescence intensity values, which are reported in **Fig. 3G** and **fig. S8C**.

### Immunoblotting

To quantify protein levels, cells were lysed in Laemmli buffer (0.0625 mM Tris-HCl, 10% glycerol, 2% SDS,5% 2-mercaptoethanol, 0.002% bromophenol blue) supplemented with RIPA buffer (50 mM Tris-HCl, 150 mM NaCl, 1% Triton-X-100, 0.1% SDS, 10 mM NaF, 0.5% sodium deoxycholate) and Halt protease and phosphatase inhibitor cocktail (ThermoScientific, 78440). Protein lysates were heated at 95 °C for 5 minutes, centrifuged at 13,000 rpm for 7 minutes at 4 °C to pellet cell debris, and the supernatant was collected for further analysis. Proteins were separated by SDS-PAGE and transferred to polyvinylidene fluoride (PVDF) membranes, which were then blocked with 3% nonfat milk and incubated overnight with primary antibodies (see **Table S4** for details). After washing with Tris-Buffered Saline + 0.05% Tween-20, membranes were incubated with HRP-conjugated IgG secondary antibodies (Santa Cruz Biotechnologies) for one hour. Protein bands were visualized using Western Lightning^TM^ Plus - Enhanced Chemiluminscence Substrate (Perkin Elmer Life & Analytical Sciences) and imaged on a ChemiDoc XRS system (Bio-Rad).

### Bulk ATP luminescence assay

We measured whole-cell ATP levels using the CellTiter-Glo Kit (Promega, G9242). ATP standard was prepared (Sigma, A7699) at concentrations from 0.01 µM to 1 µM. Cells (200 k/well) were seeded on a 6-well plate for 24 hours before drug treatment. Cells were treated with TH5427 (1 µM or 10 µM) and oligomycin A (1 µM) for 24 h. Following treatment, cells were resuspended in serum-free, phenol red-free media at 10,000 cells/mL and incubated for 1.5 h (OVCAR-5-CisR); 100 µL of cell suspension or ATP standard was added to 100 µL of CellTiter-Glo reagent. ATP levels were measured by following manufacturer’s instructions using a luminometer (SpectraMax M2, Molecular Devices).

### Adenosine-responsive aptamer sensor

For detecting adenosine metabolite levels using an aptamer sensor, siRNA-treated cells were plated at a density of 10,000 cells per cm^2^ into a 24-well plate; for NUDT5 inhibition experiments, cells were plated at a density of 5,000 cells per cm^2^, followed by treatment with 10 µM TH5427 or 1 µM oligomycin A for 24 hours. The next day, the sensor was hybridized by mixing the aptamer with the quencher strand at a 1 to 3 aptamer to quencher ratio using ATP buffer (50 mM Tris, 100 mM NaCl, 5 mM MgCl_2_ pH 7.4), heating the samples at 95 °C for 5 minutes, and cooling at room temperature for 20 minutes. For 1 well of a 24-well plate, 25 µL of the sensor was diluted in 73 µL of reduced serum Opti-MEM (ThermoFisher, 31985062), followed by the addition of Turbofect (2 µL) (ThermoScientific, R0533), and mixed immediately by pipetting up and down. While the sensor complex incubated for 20 minutes, cells were carefully washed three times with PBS and 900 µL of Opti-MEM was added to the cells. Then 100 µL of the sensor was added to the wells and gently mixed to evenly distribute the DNA. After incubating the cells with sensor for 4 hours at 37 °C, cells were washed three times with PBS. Cells were then incubated with CellTracker Red CMPTX (1 µM) (ThermoFisher, C34552) and Hoescht (1 µM) (ThermoFisher, 33342) for 15 minutes at 37 °C to label the cells and nuclei, respectively. Cells were incubated in fresh Opti-MEM for epifluorescence imaging, which was conducted using a Zeiss Axio Observer Z1 inverted fluorescence microscope equipped with a 20× objective (N.A. 0.8); for one replicate of the siRNA-treated cells with the aptamer sensor, epifluorescence imaging was conducted with a 10× objective (N.A. 0.30). To account for potentially false positive signals due to sensor degradation, cells were transfected with an inactive aptamer sensor containing a scrambled sequence modified with a 5-carboxyfluorescein (FAM) label that served as a negative control. Corresponding adenosine-responsive aptamer and quencher sequences can be found in **Table S5**.

### Targeted metabolomics

We seeded 500,000 of OVCAR-5-CisR cells in a 60 mm tissue-culture dish in culture media with 10% dialyzed FBS (dFBS). After 24 hours of cell attachment, media was replaced with drug-treated media with 10% dFBS. To minimize contributions of extracellular metabolites, media was changed 3 hours priors to extraction with drug-treated media with 10% dFBS. Each culture plate was removed from the incubator, aspirated, and quenched with 1.2 mL of a high-performance liquid chromatography (HPLC) grade solution of precooled (−20°C) 40:40:20 acetonitrile/methanol/water; extraction continued at -20 °C for 15 minutes. Cells were scraped from the plate and the entire contents were transferred into an Eppendorf tube for centrifugation 17,000 × g for 5 minutes at 4 °C. The supernatant was dried under nitrogen flow and reconstituted in 50× packed cell volume (PCV) µL of HPLC grade water for liquid chromatography-mass spectrometry (LC-MS) analysis.

The metabolite extract samples were analyzed by HPLC (Vanquish Duo UHPLC, Thermo Fisher Scientific) coupled to a high-resolution orbitrap mass spectrometer (Q Exactive Plus, Thermo Fisher Scientific). The LC separation method relied on hydrophilic interaction chromatography using an XBridge BEH Amide XP Column, 130 Å, 2.5 µm, 2.1 mm × 150 mm (Waters). MS was conducted in both the positive and negative mode using a mass resolution of 140,000 at 200 m/z. Metabolomic Analysis and Visualization Engine (MAVEN) was used to analyze the LC-MS data (*79, 80*). Peaks were identified based on known retention times and mass-to-charge ratios (m/z) (*80*).

### Patient data analysis

NUDT5 transcriptomic data associated with tumor grade, overall survival, and progression-free survival was obtained from the ovarian cancer database of Cancer Science Institute Singapore (CSIOVDB; http://csiovdb.mc.ntu.edu.tw/pages/CSIOVDB_NUDT5.html) (*48*). NUDT5 expression levels across various cancer types were obtained from The Cancer Genome Atlas (TCGA) database. Normal tissue expression levels were obtained from the Genotype-Tissue Expression (GTEx) Portal. NUDT5 expression analysis (DESeq2) by cancer types was performed using GEPIA3.

### RNAScope *in situ* hybridization (ISH)

Custom RNAscope Hs-NUDT5 20 ZZ probe targeting 1358-2434 of NM_014142.4 was designed by Bio-Techne (Catalog # 1261318-C1). RNA in situ hybridization on a tissue microarray was performed by the Translational Pathology Core Laboratory at UCLA following the manufacturer’s recommendations (ACD Bio-Techne). The slides were scanned at 40× with a digital scanner (Ariol).

### Colony formation assay

To assess *in vitro* colony formation, we seeded 1,500 OVCAR-5-CisR cells on tissue culture treated 6 well plates. After 24 h, cells were treated with 1 or 10 µM TH5427, 1 µM oligomycin A, 75 nM cytochalasin D, or vehicle (DMSO) control. Media containing each drug was replaced daily over the 10-day experimental timecourse. After 10 days, colonies were washed, fixed with 4 % paraformaldehyde for 15 minutes, and stained using 0.5% crystal violet solution (Abcam) for 30 minutes before washing with water. Colonies were allowed to dry prior to imaging. Colony images were acquired using an iPhone 13 (Apple) equipped with a 12 MP camera (ƒ/1.6 aperture) under uniform white light. To measure the colony area, we used Fiji (ImageJ) equipped with plugins ‘Colony area’ and ‘Colony measurer’.

### Mouse model experiments

All procedures in mice were performed per the NIH Guide for the Care and Use of Laboratory Animals and approved by the University of California Los Angeles Animal Research Committee (ARC-2020-011). Female C57BL6 mice (8 weeks old) were obtained from Charles River Laboratories. On day 1, mice were injected intraperitoneally with the syngeneic mouse ovarian cancer cell line SO (5 × 10^5^ cells per mouse), which forms highly aggressive tumors within 14 days (*51*). Two days after cancer cell injection, mice received a daily intraperitoneal injection of either vehicle (sterile water) or the NUDT5 inhibitor TH5427 (25 mg/kg in sterile water) for 2 weeks. On day 14, mice were euthanized, tumor burden was assessed in the omentum and at peritoneal sites beyond the omentum.

### Statistical methods

To rank compounds with statistically significant effects on cell filtration in the LOPAC screen, we calculated a Z-score for each sample, *Z = (Absorbance_Drug_ - Mean Absorbance_Vehicle_) / SD Absorbance_Vehicle_*. The cell filtration screen was performed once. Experiments were conducted over *n* independent experiments across *N* individual cells or technical replicates, as detailed in the figure captions. Data are presented as the median with 95% confidence interval (CI) or interquartile range (IQR), except where noted. Unless otherwise specified, data were analyzed using non-parametric tests: the Mann-Whitney U-test for comparisons between two groups, and Kruskal-Wallis with Dunn’s test for comparisons among three or more groups. Categorical outcomes were compared using the chi-square (χ²) test. Exact p-values are reported to two significant figures; values below 1 × 10^-15^ are reported as p < 1 × 10^-15^. We considered p-values less than 0.05 to be statistically significant. GraphPad Prism 10.0 or 9.0 software was used for all statistical evaluations.

## Acknowledgments

Funding

Research was supported by Allen Family Philanthropies Distinguished Investigator Awards (ACR, DNR, YL); the Department of Defense Ovarian Cancer Research Program TEAL Expansion Award (ACR, SO, BYK); the National Science Foundation BRITE Fellow Award grant CMMI-2135747 (ACR); the National Institutes of Health (NIH) grant R35GM161448 (ACR), R01NS130677 (SL), and R35GM141931 (YL), and R35GM143127 (JOP); the National Cancer Institute grant R21CA245667 (ACR, SO, RD, BYK); a National Cancer Institute/ NIH Fellowship F31CA288105, Whitcome Fellowship, and UCLA Jonsson Comprehensive Cancer Center Fellowship (AMF).

## Author contributions

ACR, RD, SO, BYK, NKG, AMF, and JS conceptualized the project; AMF, JS, NKG, AA, JOP, MSR, DQ, CL, RK, EP conducted experiments; VG and YL generated and provided the adenosine-responsive aptamer; ACR, RD, SO, SL, DNR, JOP, and YL provided supervision; SO conducted microarray data analysis; AMF, SO performed transcriptomic connectivity analysis; AMF, JS, NKG, AA, EP, MSR, SO, ACR contributed to writing – methods; AMF, JS, MSR, SO, ACR contributed to writing – results; ACR wrote and refined the manuscript; all reviewed and edited the final manuscript.

## Data and materials availability

All data are available in the main text or the supplementary materials. RNA sequencing data generated in this study have been deposited in the NCBI Gene Expression Omnibus (GEO) under accession number GSE335942. Code is freely available through github. Additional data related to this paper may be requested from the authors. We are grateful to Amanda Faye Lipsey, of Amanda Faye Consulting, for her support in preparing this manuscript.

