## Supplementary Figures and Tables for "Deformability screening identifies NUDT5 as a mediator of cellular mechanobiology"

This PDF file includes:

- Supplemental Figures S1 to S14
- Supplemental Tables S1 to S5

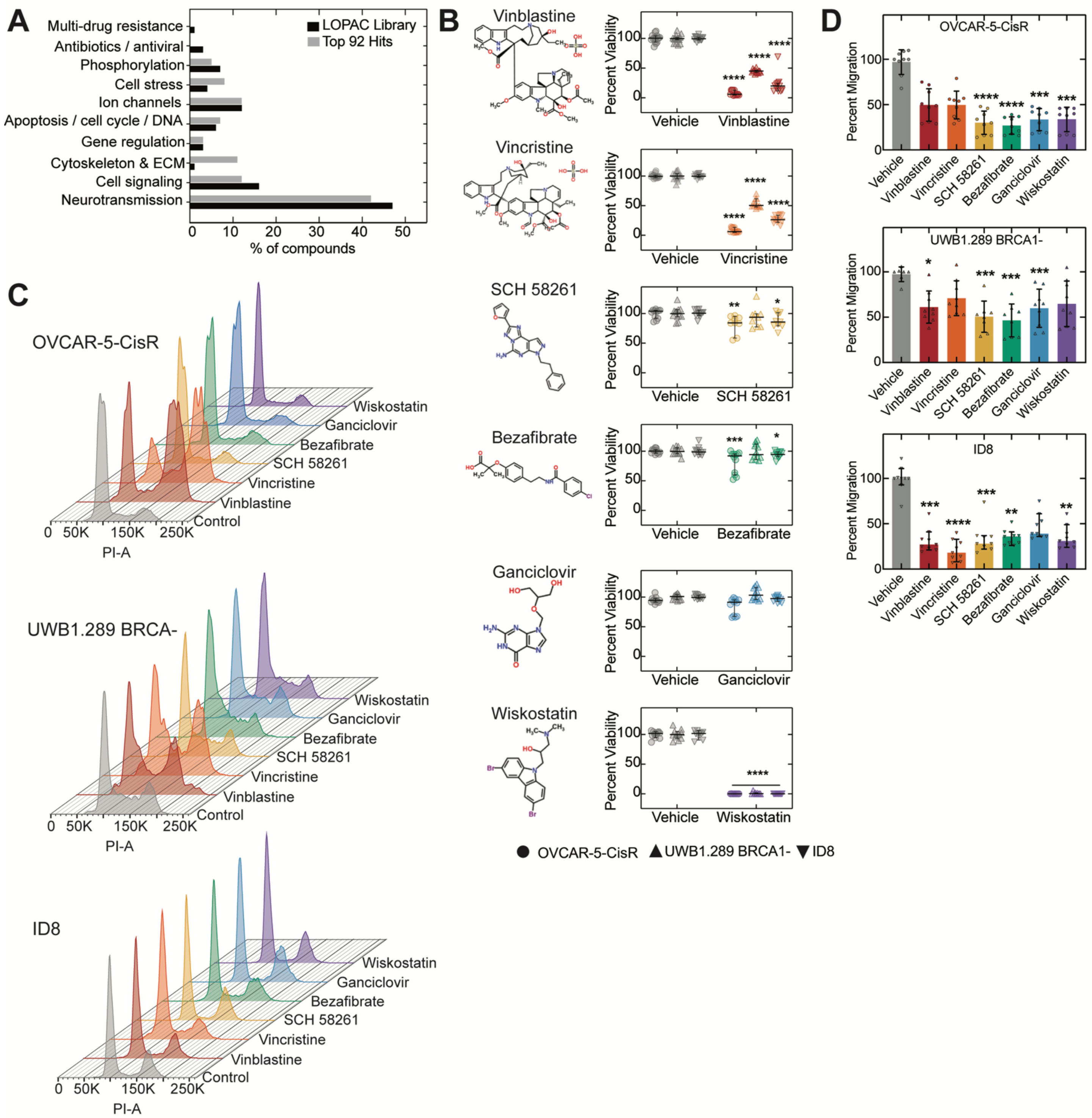


**Fig. S1. Secondary assays to validate small molecules identified from deformability screen. (A)** Distribution across drug categories for the 1280 compounds of the LOPAC library (Sigma) and the top 92 hits with Z > 2. (**B**) Chemical structures of the top six compounds identified in the deformability screen and their effects on cell viability measured by CellTiter-Glo assay across cell lines (OVCAR-5-CisR, UWB1.BRCA-1, ID8). The percentage of viable cells is determined relative to vehicle control (24 h treatment). (**C**) Representative data showing the distributions of propidium iodide area (PI-A) signal to determine cell cycle distributions after 48 h drug treatments using flow cytometry. Shown here are data for OVCAR-5-CisR, UWB.289 BRCA, and ID8 cells treated with same concentrations of drugs as in S1.A; quantification of cell cycle distributions across n =2 independent experiments are shown in **Figure 1H**. (**D**) Migration is measured after 24 h of a transwell assay (24 h treatment). We quantified the number of migrated cells normalized to the vehicle (percentage) through a membrane with 8 µm pores after 24 h. All data in this figure shows results for 0.5 nM for vinblastine and vincristine; 0.1 µM for wiskostatin; and 1 µM for SCH 58261, bezafibrate, and ganciclovir. Bars show median; errors bars represent 95% confidence interval. Data shown for n =3 independent experiments. Statistical significance was determined using a Mann-Whitney test for (**D**), *p < 0.05, **p < 0.01, ***p < 0.001, ****p < 0.0001.

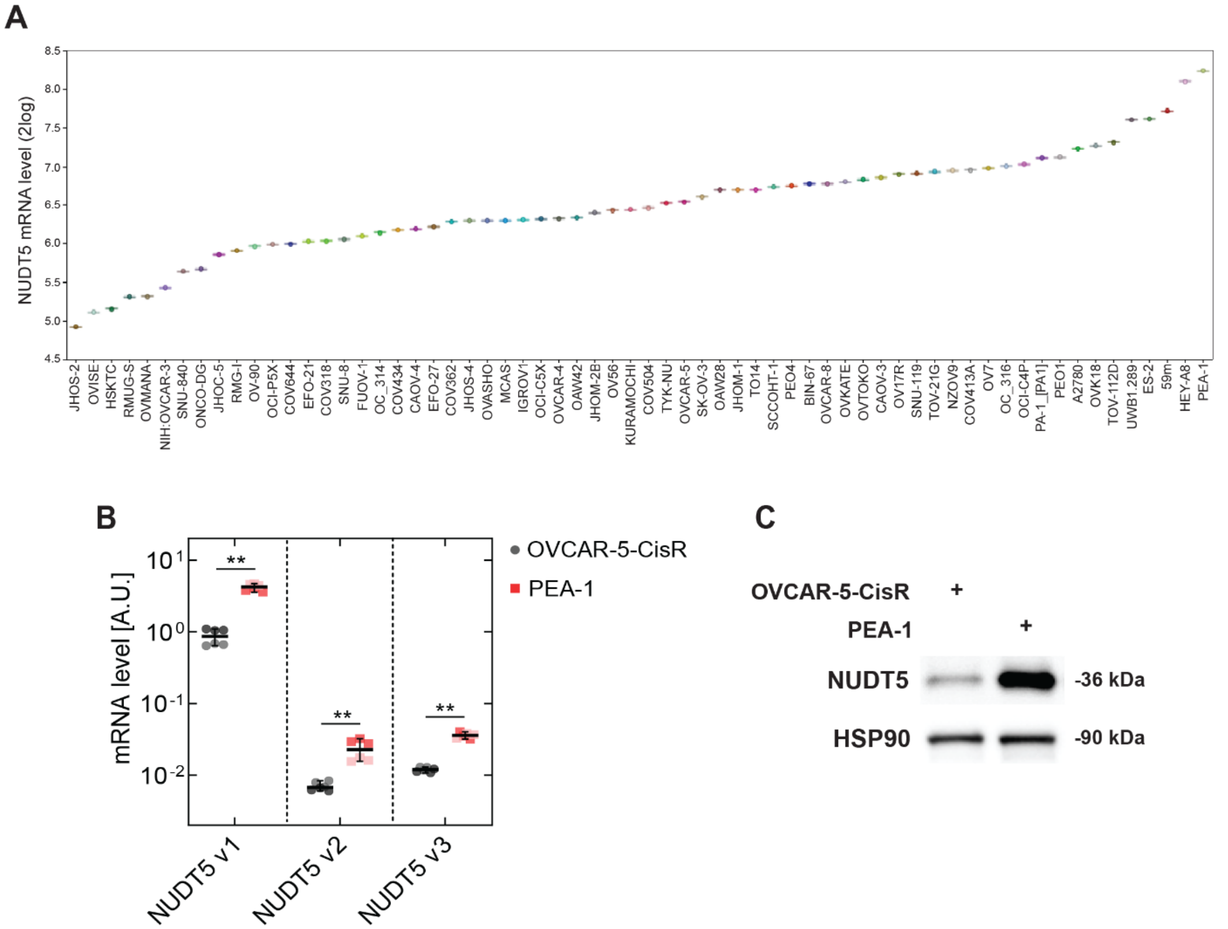


**Fig. S2. NUDT5 expression in various human ovarian cancer cell lines.** (**A**) mRNA levels of NUDT5 in human ovarian cancer cell lines extracted from the Cancer Cell Line Encyclopedia (CCLE) 21q4 (N = 64 cell lines). (**B**) mRNA levels of NUDT5 isoforms in OVCAR-5-CisR and PEA-1 cells, as determined using qRT-PCR. Each dot represents a technical replicate across two independent experiments (N = 6 replicates for OVCAR-5-CisR and PEA-1); the horizontal line represents the median with 95% confidence interval. Statistical significance was determined using a Mann-Whitney test (**$p < 0.01$). (**C**) Immunoblot of NUDT5 in OVCAR-5-CisR and PEA-1 cells, where heat shock protein 90 (HSP90) serves as a loading control (n = 1).

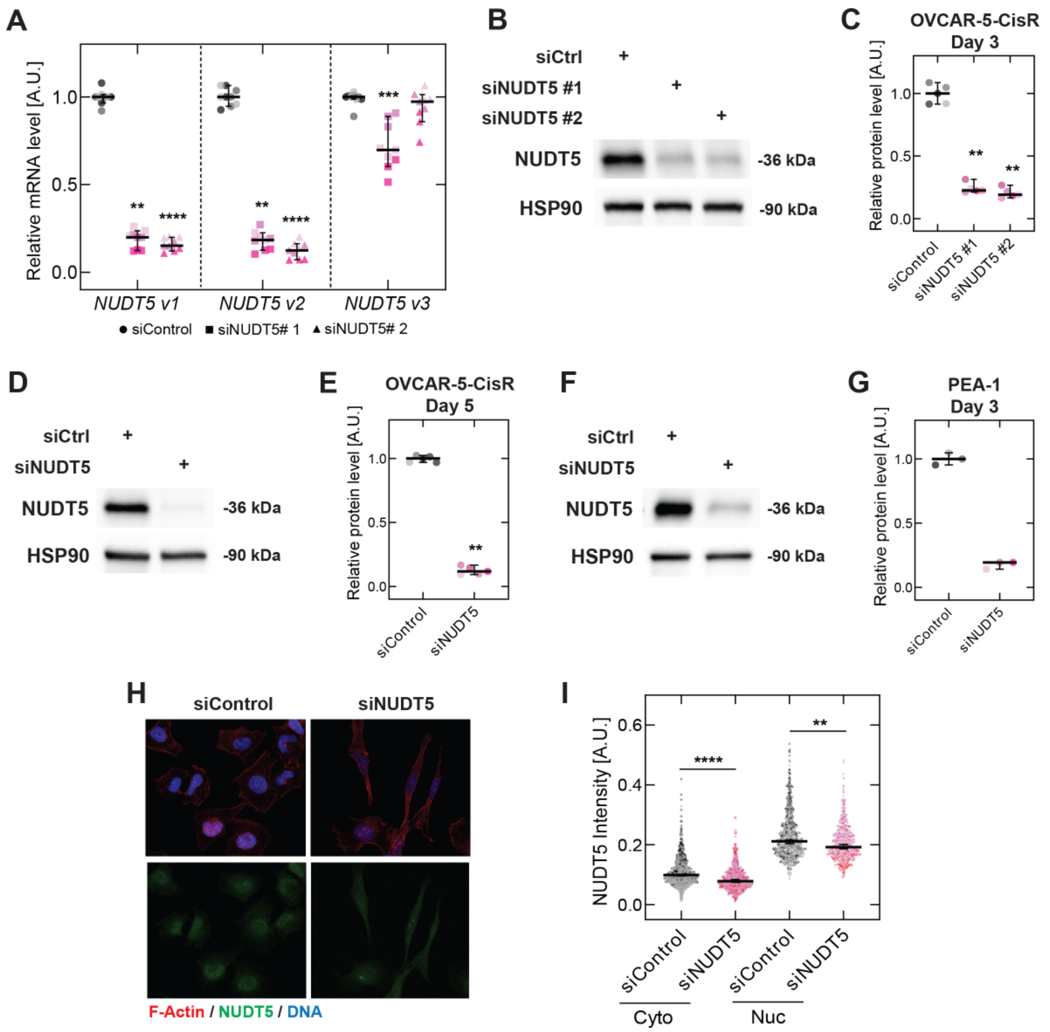


**Fig. S3. Characterization of NUDT5 levels with siNUDT5 knockdown in OVCAR-5-CisR and PEA-1 cells.** (**A**) mRNA levels of NUDT5 isoforms in OVCAR-5-CisR cells treated with siControl and siNUDT5 on day 3 post-transfection, as determined using qRT-PCR and normalized to siControl (N = 9 replicates). (**B**) Representative immunoblot of NUDT5 in OVCAR-5-CisR cells on day 3 after siNUDT5 knockdown, where HSP90 serves as a loading control. (**C**) Quantification of protein levels in OVCAR-5-CisR cells on day 3 normalized to siControl (n = 5 independent experiments). (**D**) Representative immunoblot of NUDT5 in OVCAR-5-CisR cells on day 5 after siNUDT5 knockdown, where HSP90 serves as a loading control. (**E**) Quantification of protein levels in OVCAR-5-CisR cells on day 5 normalized to siControl (n = 5 independent experiments). (**F**) Representative immunoblot of NUDT5 in PEA-1 cells on day 3 after siNUDT5 knockdown, where HSP90 serves as a loading control. (**G**) Quantification of protein levels in PEA-1 cells on day 3 normalized to siControl (n = 3 independent experiments). (**H**) Confocal images of F-actin (phalloidin, red), DNA (DAPI, blue), and NUDT5 (green) in siControl and siNUDT5-treated OVCAR-5-CisR cells on day 3. Scale, 25 µm. (**I**) Quantification of NUDT5 intensity in cytoplasmic and nuclear regions based on images in **H**. In **A**, **C**, **E**, **G**, and **I**, scatter plots show median values with 95% confidence interval, where each dot represents a single replicate (**A**, **C**, **E**, **G**) or cell/nuclei (**I**) across n = 3-5 independent experiments. In **A** and **I**, statistical significance was determined by a Kruskal-Wallis with Dunn's multiple comparison test (**p < 0.01, ***p < 0.001, ****p < 0.0001). In **C** and **E**, statistical significance was determined using a Mann-Whitney test compared to the control (**p < 0.01).

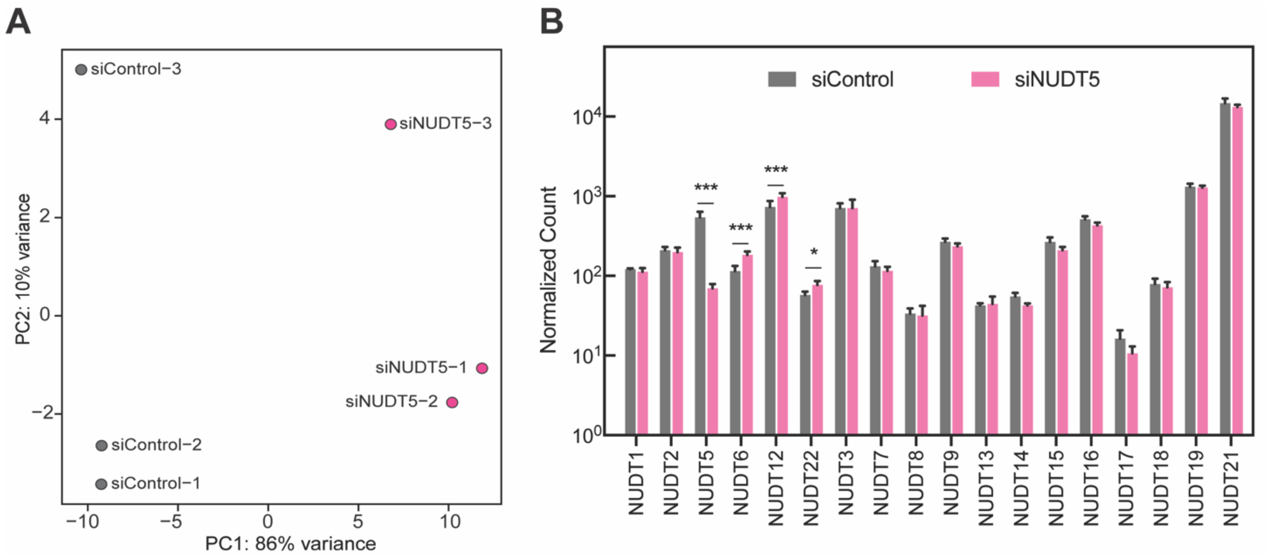


**Fig. S4. Evaluating OVCAR-5-CisR cells with NUDT5 knockdown (siNUDT5) using bulk RNA sequencing (RNAseq).** (**A**) Principal component analysis (PCA) of bulk RNAseq data from n = 3 independent biological replicates of siNUDT5 and siControl OVCAR-5-CisR cells on day 2 post-transfection. (**B**) DESeq2-normalized mRNA counts for individual NUDT gene family members in siNUDT5 and siControl OVCAR-5-CisR cells. Bar represents mean of n = 3 independent biological replicates; error bars represent standard deviation. Significant differentially expressed genes were determined using a cutoff of padj < 0.05 from DESeq2 analysis (*p < 0.05, ***p < 0.001).

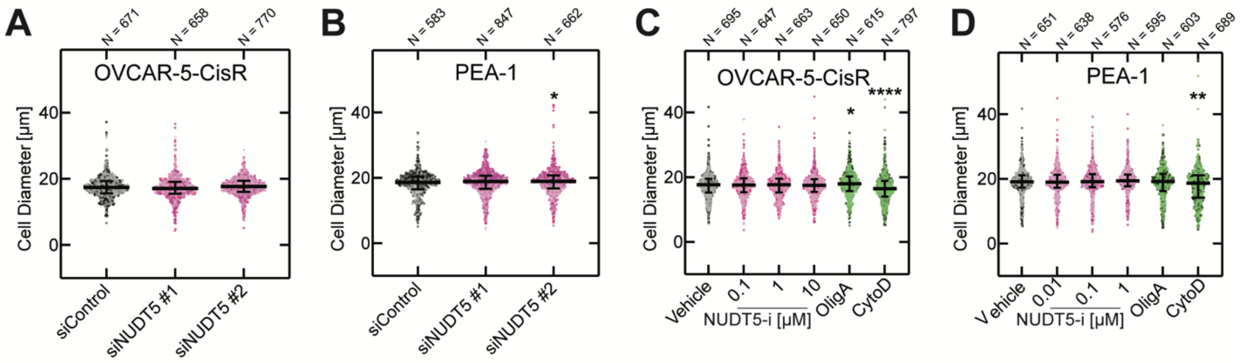


**Fig. S5. Characterization of cell size for OVCAR-5-CisR and PEA-1 cells with NUDT5 manipulations.** Measurements of the diameter of cells in a suspended state prior to filtration using quantitative image analysis for (**A**) OVCAR-5-CisR and (**B**) PEA-1 cells treated with siControl and siNUDT5. Measurements of cell diameter for (**C**) OVCAR-5-CisR and (**D**) PEA-1 cells treated with vehicle (DMSO), NUDT5-i (TH5427), oligomycin A (ATP synthase-I or OligA) or cytochalasin D (CytoD) for 24 hours. Each dot represents an individual cell; horizontal lines show median values; error bars represent 95% confidence interval. Data shown is acquired over n = 3 independent experiments. Statistical significance was determined by a Kruskal-Wallis with Dunn's multiple comparison test compared to vehicle control (*p < 0.05, **p <0.005, ****p < 0.0001).

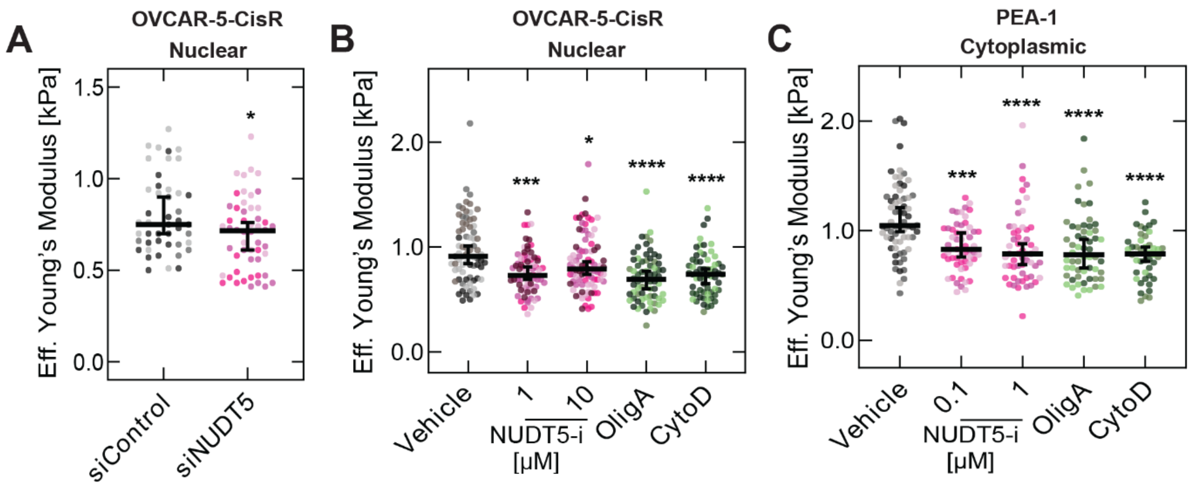


**Fig. S6. Characterizing the stiffness of cellular and nuclear regions using force probe indentation.** (**A**) Effective Young's modulus of the nuclear region of OVCAR-5-CisR cells treated with siControl and siNUDT5 on day 3 post-transfection, as determined using force probe indentation (N = 45 cells for siControl, N = 52 cells for siNUDT5). (**B**) Effective Young's modulus of the nuclear region of OVCAR-5-CisR cells treated with vehicle (N = 73 cells), NUDT5 inhibitor (TH5427) (N = 74 cells for 1 μM, N = 85 cells for 10 μM), Oligomycin A (N = 67 cells) or cytochalasin D (N = 62 cells) for 24 hours. (**C**) Effective Young's modulus of the cytoplasmic region of PEA-1 cells treated with vehicle (N = 66 cells), NUDT5 inhibitor (TH5427) (N = 55 cells for 0.1 μM, N = 56 cells for 1 μM), Oligomycin A (N = 58 cells) or cytochalasin D (N = 44 cells) for 24 hours. Plots show median values with 95% confidence interval, where each dot represents a single cell across n = 3 independent experiments. In **A,** statistical significance was determined using a Mann-Whitney test compared to the control (*p < 0.05). In **B** and **C**, statistical significance was determined by a Kruskal-Wallis with Dunn's multiple comparison test where each treatment was compared to the vehicle (*p < 0.05, ***p <0.001, ****p < 0.0001).

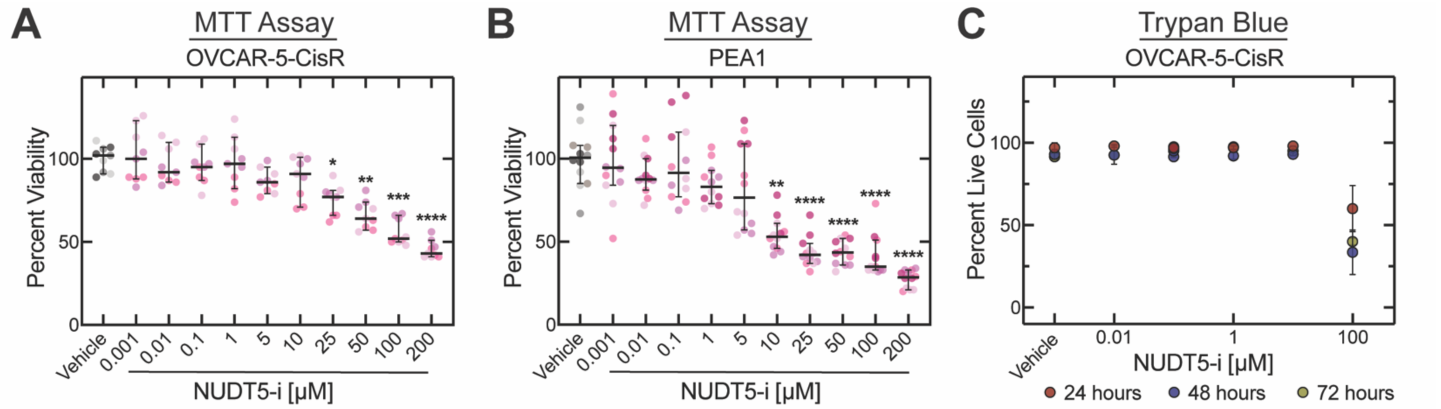


**Fig. S7. Determination of the viability of cells treated with the NUDT5 inhibitor (NUDT5-i), TH5427.** (**A-B**) Cell viability of OVCAR-5-CisR (**A**) and PEA-1 (**B**) cells after treatment with varying doses of NUDT5-i (TH5427) for 24 h, as determined using an MTT assay (n = 3 independent experiments for OVCAR-5-CisR; n = 4 for PEA-1 cells). Data show the percentage of viable cells relative to vehicle control. (**C**) Percentage of viable OVCAR-5-CisR cells after treatment with varying doses of NUDT5-i (TH5427) for 24 - 72 h, as determined using a Trypan blue assay (n = 2 independent experiments). Plots show median values with 95% confidence interval. Statistical significance was determined by a Kruskal-Wallis with Dunn's multiple comparison test compared to vehicle control (*p < 0.05, **p <0.01, ***p < 0.005, ****p < 0.0001).

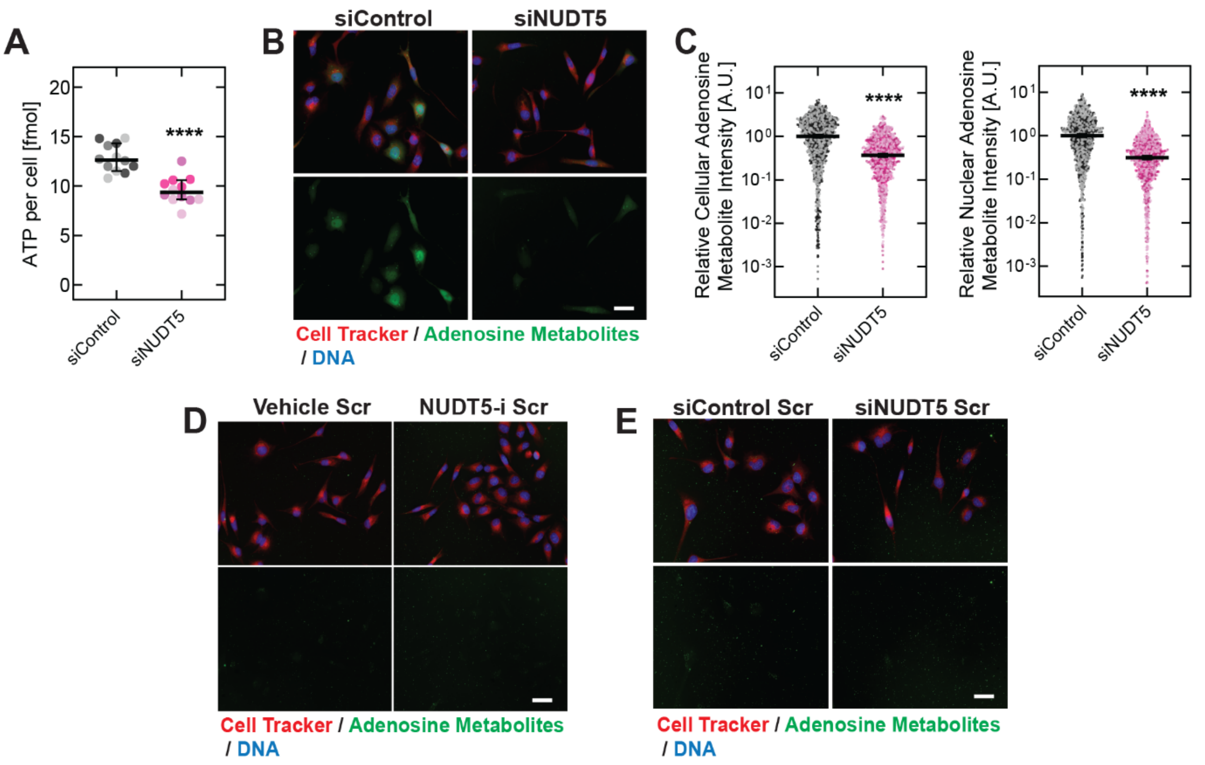


**Fig. S8. NUDT5 knockdown reduces ATP levels in OVCAR-5-CisR cells.** (**A**) Cellular ATP levels of cells treated with siControl and siNUDT5, as measured using a luminescence-based ATP assay (N = 12 replicates). We assayed cells on day 2 post-transfection. (**B**) Epifluorescence images of an adenosine-responsive aptamer (green) in siControl and siNUDT5-treated cells on day 3 post-transfection. Cells are visualized using CellTracker (red) and DAPI (blue). Scale, 50 µm. (**C**) Quantification of adenosine metabolite levels across the whole cell or nuclear region based on images in **B** (N > 950 cells for siControl and siNUDT5). (**D**) Epifluorescence images of a scrambled non-responsive aptamer (green) in vehicle (DMSO) or NUDT5-i (TH5427)-treated cells. Scale, 50 µm. (**E**) Epifluorescence images of a scrambled non-responsive adenosine aptamer (green) in siControl and siNUDT5-treated cells on day 3 post-transfection. Scale, 50 µm. In **A** and **C**, scatter plots show median values with 95% confidence interval, where each dot represents a technical replicate (**A**) or individual cell/nucleus (**C**) across n = 3 independent experiments. Statistical significance was determined using a Mann-Whitney test (****$p < 0.0001$).

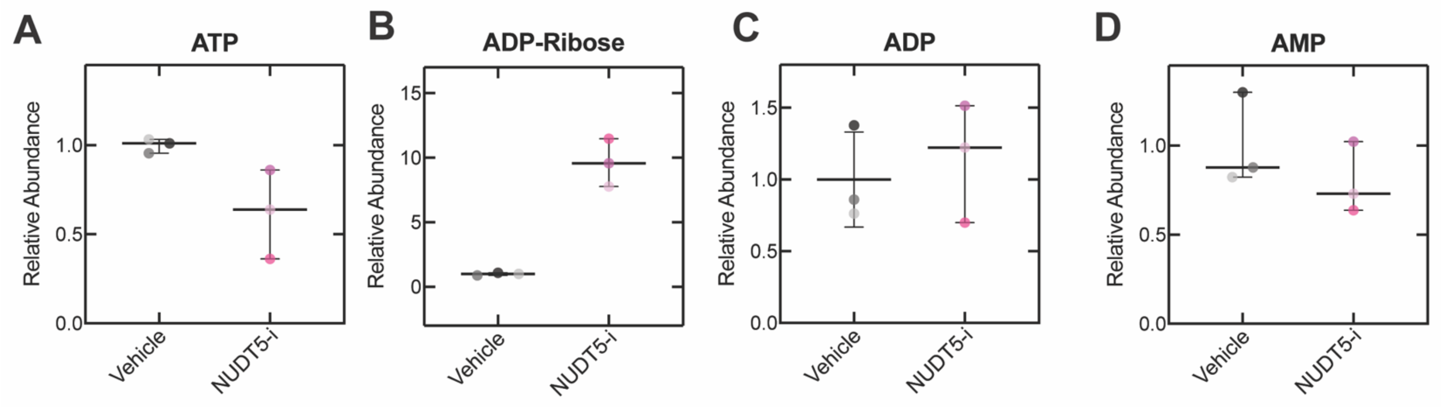


**Fig. S9. Targeted metabolomics of OVCAR-5-CisR cells with NUDT5 inhibition.** Cells were treated with vehicle (DMSO) or 10 µM NUDT5-i (TH5427) for 24 h. Abundance of ATP ($p = 0.1$) (**A**), ADP-Ribose ($p = 0.1$) (**B**), ADP ($p > 1.0$) (**C**), and AMP ($p = 0.4$) (**D**) were measured and normalized to the vehicle. Data show median across $n = 3$ independent experiments represented by individual symbols; error bars represent 95% confidence interval.

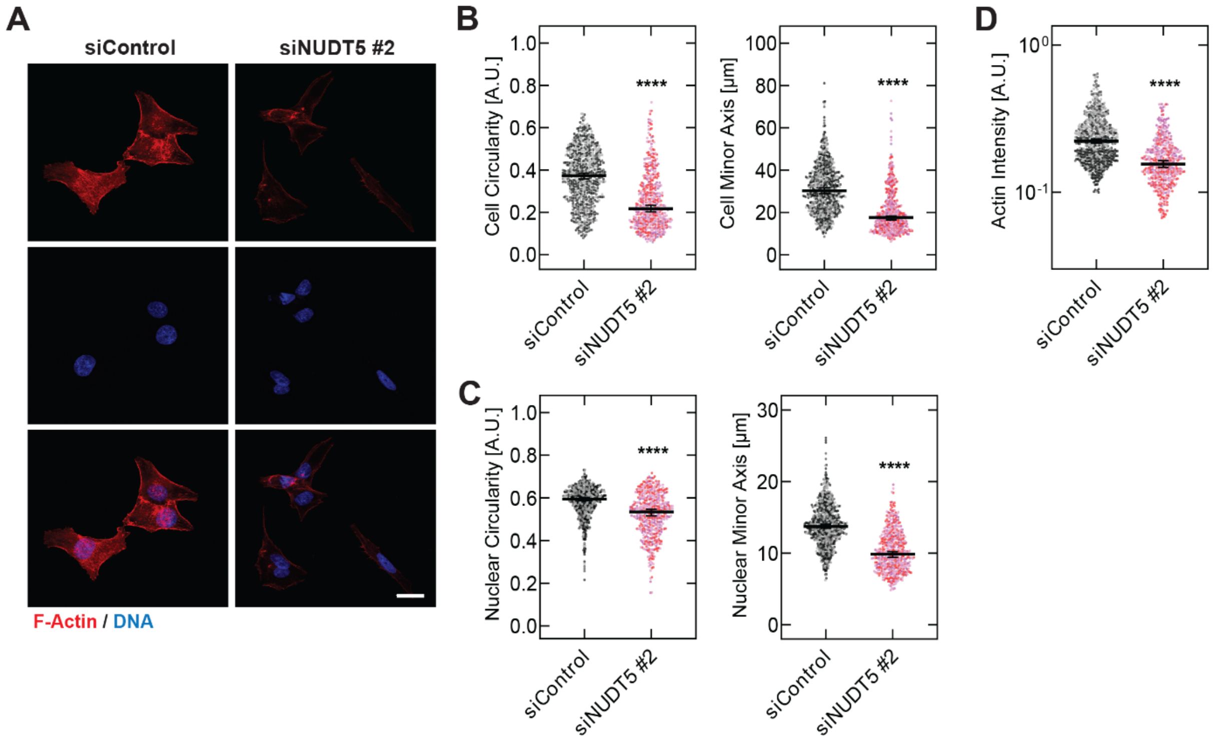


**Fig. S10. Effects of NUDT5 knockdown on OVCAR-5-CisR cell morphology.** (**A**) Confocal images of F-actin (phalloidin, red) and DNA (DAPI, blue) in OVCAR-5-CisR cells treated with siControl and siNUDT5 on day 3 post-transfection. Scale, 25 µm. (**B**-**D**) Quantitative image analysis for measurements of (**B**) cell circularity, cell minor axis length; (**C**) nuclear circularity; nuclear minor axis length; and (**D**) F-actin intensity. For cell circularity, minor axis length and F-actin intensity, data shown represent N = 678 cells for siControl, N = 475 cells for siNUDT5; for nuclear circularity and minor axis length: N = 674 cells for siControl, N = 457 cells for siNUDT5. Scatter plots show median values with 95% confidence interval, where each dot represents a single cell or nucleus across n = 3 independent experiments. Statistical significance was determined using a Mann-Whitney test (****$p < 0.0001$).

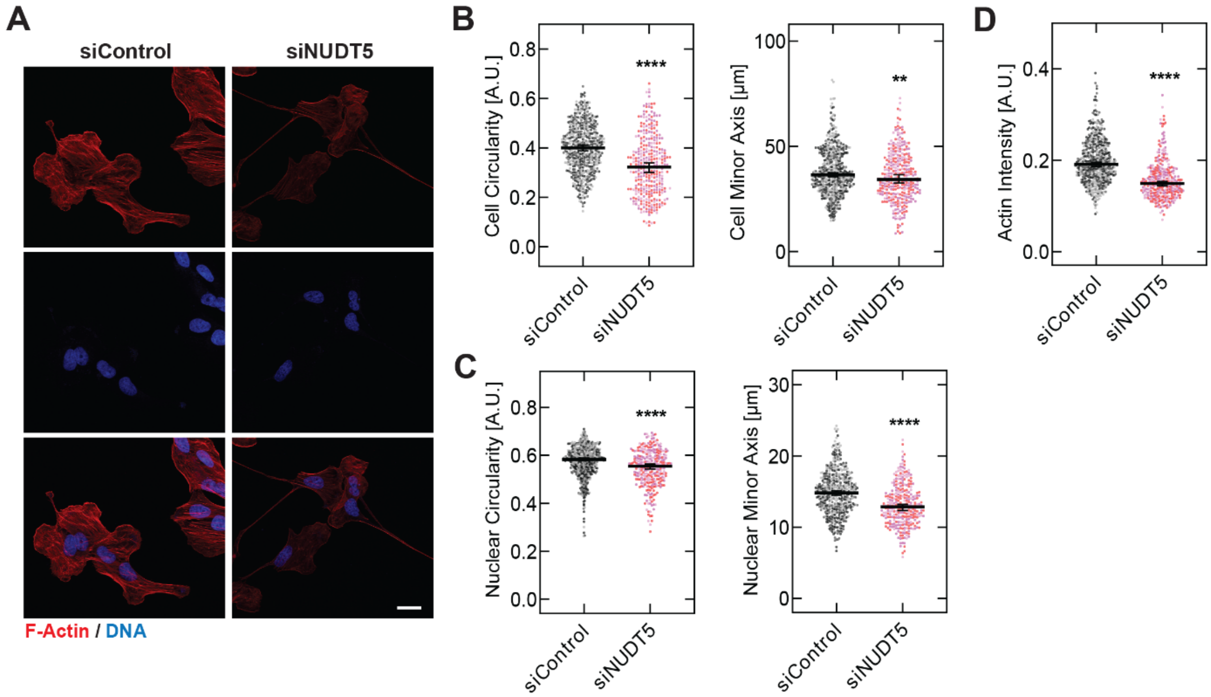


**Fig. S11. Effects of NUDT5 knockdown on PEA-1 cell morphology.** (**A**) Confocal images of F-actin (phalloidin, red) and DNA (DAPI, blue) in PEA-1 cells treated with siControl and siNUDT5 on day 3 post-transfection. Scale, 25 µm. (**B**-**D**) Quantitative image analysis for measurements of (**B**) cell circularity, cell minor axis length; (**C**) nuclear circularity; nuclear minor axis length; and (**D**) F-actin intensity. For cell circularity, minor axis length and F-actin intensity, data represent N = 515 cells for siControl, N = 324 cells for siNUDT5; for nuclear circularity and minor axis length: N = 486 cells for siControl, N = 293 cells for siNUDT5. Scatter plots show median values with 95% confidence interval, where each dot represents a single cell across n = 3 independent experiments. Statistical significance was determined using a Mann-Whitney test (**$p < 0.01$, ****$p < 0.0001$).

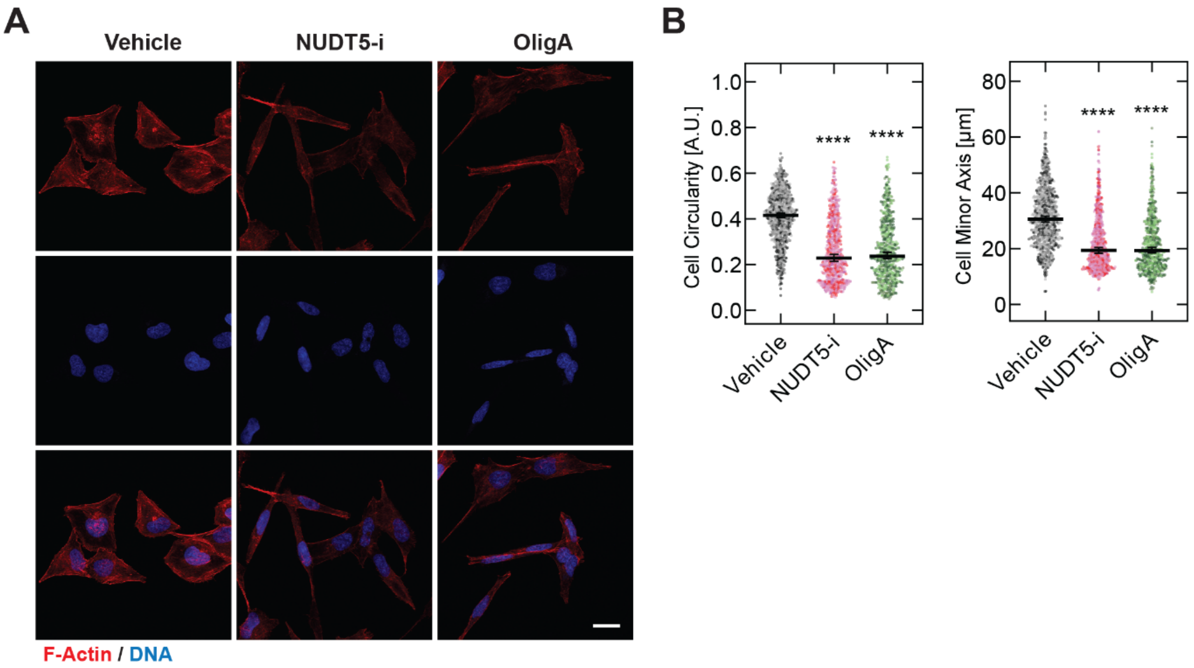


**Fig. S12. Effects of NUDT5 inhibition on OVCAR-5-CisR cell morphology.** (**A**) Confocal images of F-actin (phalloidin, red) and DNA (DAPI, blue) in OVCAR-5-CisR cells treated with vehicle, NUDT5 inhibitor (TH5427) or Oligomycin A for 24 hours. Scale, 25 µm. (**B**) Quantitative image analysis for measurements of cell circularity and cell minor axis length. Data shown represent N = 1071 cells for vehicle; N = 665 cells for NUDT5 inhibitor; N = 583 cells for OligA. Scatter plots show median values with 95% confidence interval, where each dot represents a single cell across n = 3 independent experiments. Statistical significance was determined by a Kruskal-Wallis with Dunn's multiple comparison test where each treatment was compared to the vehicle (****$p < 0.0001$).

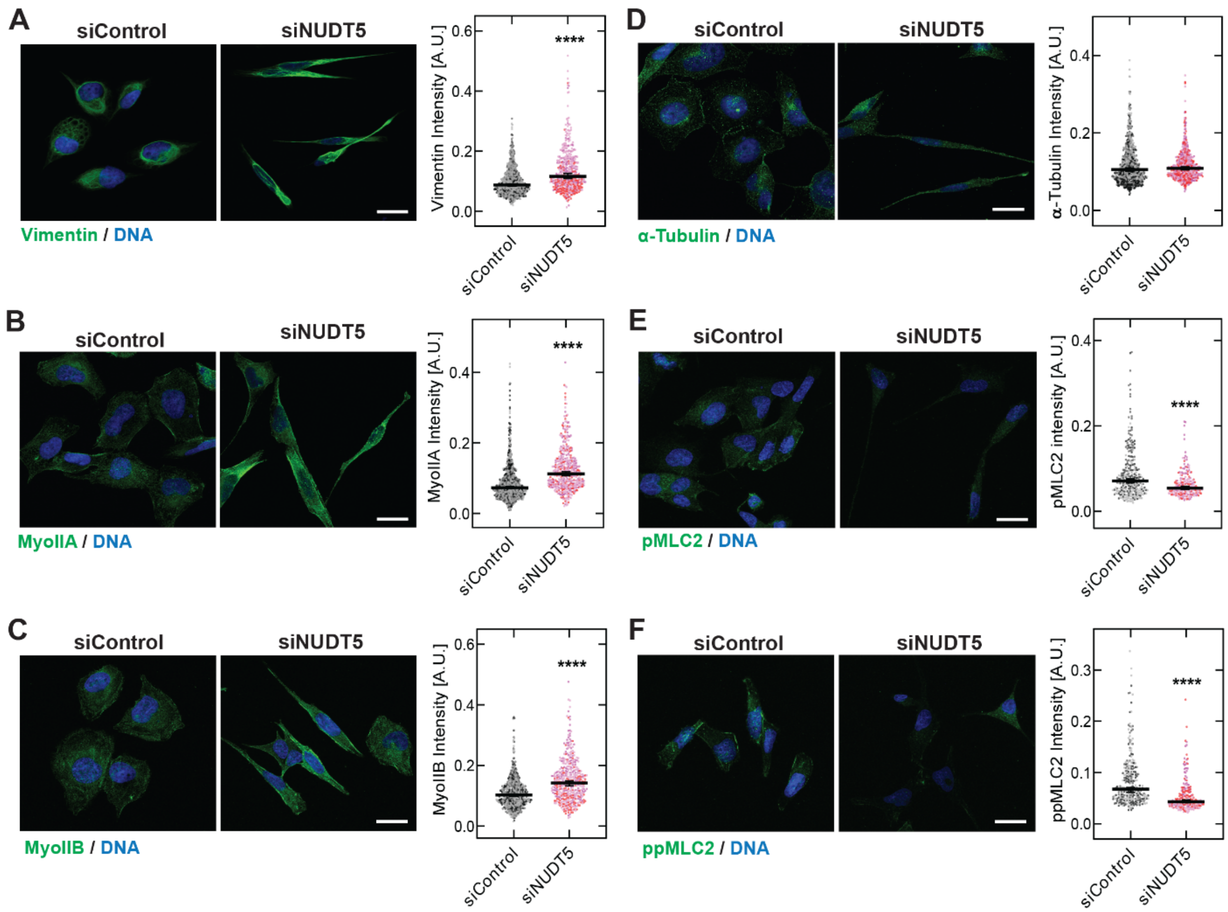


**Fig. S13. Effects of NUDT5 knockdown on cytoskeletal proteins.** (**A**-**F**) Confocal images of immunolabeled vimentin (**A**), non-muscle myosin II A (MyoIIA) (**B**), non-muscle myosin II B (MyoIIB) (**C**), α-tubulin (**D**), phosphorylated myosin light chain 2 (pMLC2) (**E**), diphosphorylated MLC2 (ppMLC2) (**F**) and DNA (DAPI, blue) in OVCAR-5-CisR cells treated with siControl and siNUDT5 on day 3 post-transfection. Scale, 25 µm. Quantification of fluorescence intensity based on corresponding images in **A**-**F** (Vimentin: N = 838 cells for siControl, N = 424 cells for siNUDT5; MyoIIA: N = 846 cells for siControl, N = 457 cells for siNUDT5; MyoIIB: N = 690 cells for siControl, N = 425 cells for siNUDT5; α-tubulin: N = 750 cells for siControl, N = 502 cells for siNUDT5; pMLC2: N = 285 cells for siControl, N = 202 cells for siNUDT5; ppMLC2: N = 273 cells for siControl, N = 216 cells for siNUDT5). Scatter plots show median values with 95% confidence interval, where each dot represents a single cell across n = 3 independent experiments. Statistical significance was determined using a Mann-Whitney test (****$p < 0.0001$).

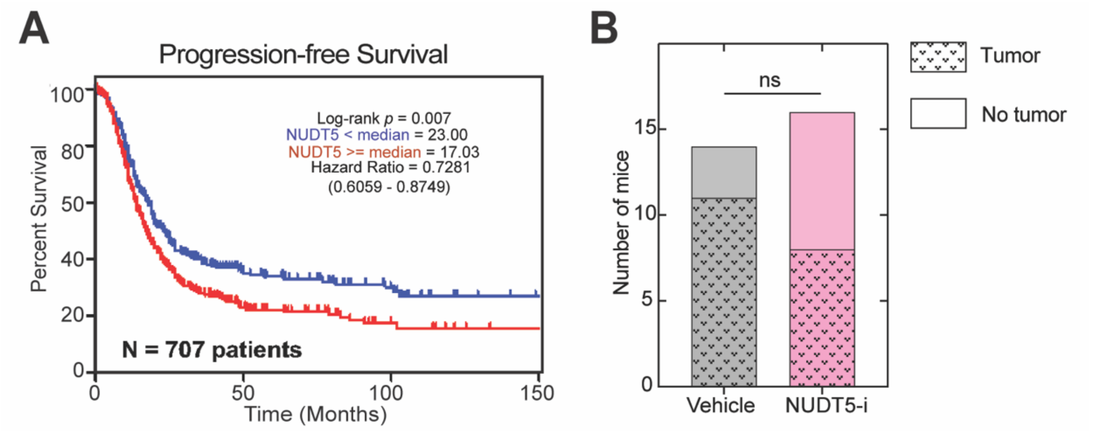


**Fig. S14. Functional relevance of NUDT5 levels and NUDT5 inhibition *in vivo*.** (**A**) Levels of NUDT5 transcripts in epithelial ovarian carcinomas are associated with progression-free survival. Data from CSIOVDB: A microarray gene expression database of ovarian cancer subtypes (*48*). (**B**) Quantification of mice with or without tumors in the omentum after treatment with NUDT5-i (TH5427) for 14 days. N = 14 mice for vehicle (sterile water) and N = 16 mice for NUDT5-i treatment over n = 3 independent experiments. Statistical significance was assessed using a Chi-Square ($\chi^2$) test (ns = not significant).

**Table S1. 92 drug hits identified from the deformability-based screen.** Results from the cellular filtration assay, including Z-scores and retention volume, are reported along with the manufacturer-annotated drug class for each compound. Drug class annotations were used for the enrichment analyses shown in **Fig. 1D-E**. Related classes were grouped into broader categories for the log2 fold-change analysis shown in **Fig. 1C**.

**Table S2.** Target sequences for siNUDT5 #1 (Hs_NUDT5_7 from Qiagen) and #2 as referenced by Page et al (Integrative DNA Technologies) (*36*). Target sequence for siControl was not provided by the manufacturer.

**Table S3. Primers used for qRT-PCR analysis.**

**Table S4. Antibody information for immunofluorescence and Western blotting analysis**

**Table S5. Adenosine aptamer and quencher sequences**

**Table S1.** 92 drug hits identified from the deformability-based screen.

| Name | Class | Z-score | Retention Volume [μL] |
|---|---|---|---|
| Vinblastine sulfate salt | Cytoskeleton and ECM | 5.07 | 353.18 |
| Vincristine sulfate | Cytoskeleton and ECM | 4.96 | 347.39 |
| SCH 58261 | Purinoceptor | 4.51 | 324.62 |
| Bezafibrate | Transcription | 4.46 | 322.2 |
| Ganciclovir | Cell Cycle | 4.42 | 320.03 |
| Wiskostatin | Actin | 4.41 | 319.67 |
| 5HPP-33 | Cytoskeleton and ECM | 4.4 | 318.99 |
| S-(-)-Eticlopride hydrochloride | Dopamine | 4.34 | 316.33 |
| Zimelidine dihydrochloride | Serotonin | 4.23 | 310.78 |
| CID2858522 | Gene Regulation | 4.22 | 310.04 |
| Felodipine | Ca2+ Channel | 4.17 | 307.72 |
| Sunitinib malate | Tyrosine Kinase | 4.08 | 302.83 |
| Fenspiride hydrochloride | Adrenoceptor | 4 | 298.95 |
| Rofecoxib | Lipid Signaling | 3.94 | 295.59 |
| Epinastine hydrochloride | Histamine | 3.82 | 289.52 |
| N-Acetyl-L-Cysteine | Glutamate | 3.76 | 286.68 |
| Genipin | Cell Biology | 3.75 | 286.11 |
| NBI 27914 | Neurotransmission | 3.69 | 283.02 |
| Tamoxifen | Antibiotics/Phosphorylation | 3.68 | 282.41 |
| Urapidil, 5-Methyl- | Adrenoceptor | 3.65 | 281.21 |
| SKF-89145 hydrobromide | Dopamine | 3.65 | 280.9 |
| S-Ethylisothiourea hydrobromide | Nitric Oxide | 3.57 | 277.21 |
| Ropinirole hydrochloride | Dopamine | 3.57 | 277.11 |
| Furegrelate sodium | Phosphorylation | 3.53 | 275.01 |
| CP-226269 | Dopamine | 3.46 | 271.67 |
| Fexofenadine hydrochloride | Histamine | 3.34 | 265.29 |
| NS5806 | Ion Channels | 3.33 | 265.05 |
| Ellipticine | Cell Cycle | 3.28 | 262.49 |
| PHA 767491 hydrochloride | kinase phosphotase | 3.18 | 257.02 |
| EGTA | Biochemistry | 3.17 | 256.73 |
| Fusaric acid | Dopamine | 3.1 | 253.43 |
| PD-180970 | Cell signaling/ tyrosine kinase | 3.06 | 251.35 |
| N-(4-Amino-2-chlorophenyl)phthalimide | Anticonvulsant | 3.06 | 251.17 |
| Alprenolol hydrochloride | Adrenoceptor | 3.05 | 250.89 |
| Rilmenidine hemifumarate | Imidazoline | 3.02 | 249.21 |
| Apomorphine hydrochloride hemihydrate | Dopamine | 2.93 | 244.54 |
| (±)-Chloro-APB hydrobromide | Dopamine | 2.87 | 241.4 |
| (±)-Atenolol | Adrenoceptor | 2.86 | 241.06 |
| 1-Phenyl-3-(2-thiazolyl)-2-thiourea | Dopamine | 2.84 | 240.04 |
| Nocodazole | Cytoskeleton and ECM | 2.84 | 239.83 |
| Roslin 2 | Cytoskeleton and extracellular matrix | 2.83 | 239.67 |
| Cinnarizine | Ca2+ Channel | 2.83 | 239.25 |
| WB-4101 hydrochloride | Adrenoceptor | 2.81 | 238.58 |
| L-741,626 | Dopamine | 2.76 | 235.72 |
| Dihydroouabain | Ion Pump | 2.72 | 234.08 |
| Nimustine hydrochloride | DNA | 2.72 | 233.88 |
| Colchicine | Cytoskeleton and ECM | 2.71 | 233.32 |
| Pentoxifylline | Cyclic Nucleotides | 2.69 | 232.41 |
| Mibefradil dihydrochloride | Ca2+ Channel | 2.69 | 232.26 |
| Cirazoline hydrochloride | Adrenoceptor | 2.68 | 231.73 |
| N-Methyldopamine hydrochloride | Dopamine | 2.66 | 230.87 |
| Amiodarone hydrochloride | Adrenoceptor | 2.66 | 230.71 |
| Oleic Acid | Phosphorylation | 2.65 | 230.3 |
| Mexiletene hydrochloride | Na+ Channel | 2.65 | 230.19 |
| L-azetidine-2-carboxylic acid | Biochemistry | 2.64 | 229.81 |
| Furafylline | Biochemistry | 2.63 | 229.52 |
| WAY-100635 maleate | Serotonin | 2.59 | 227.42 |
| Nomifensine maleate | Dopamine | 2.58 | 226.56 |
| Ro 25-6981 hydrochloride | Glutamate | 2.56 | 225.77 |
| 1-Methylnicotinamide chloride | Inflammation | 2.51 | 223.48 |
| 2-Methylthioadenosine triphosphate tetrasodium | P2 Receptor | 2.48 | 221.49 |
| NO-711 hydrochloride | GABA | 2.44 | 219.78 |
| (±)-Chlorpheniramine maleate | Histamine | 2.42 | 218.68 |
| Amifostine | Cell Stress | 2.36 | 215.58 |
| Chlorprothixene hydrochloride | Dopamine | 2.36 | 215.41 |
| Amlexanox | Histamine | 2.35 | 215.18 |
| L-Cysteinesulfinic Acid | Glutamate | 2.32 | 213.59 |
| NNC 55-0396 | Ca2+ Channel | 2.28 | 211.66 |
| D-Cycloserine | Glutamate | 2.26 | 210.67 |
| 2,3-Dimethoxy-1,4-naphthoquinone | Cell Stress | 2.26 | 210.37 |
| Atropine sulfate | Cholinergic | 2.26 | 210.31 |
| N,N,N',N'-Tetramethylazodicarboxamide | Cell Stress | 2.24 | 209.34 |
| Capecitabine | Gene Regulation | 2.23 | 208.82 |
| (-)-Quinpirole hydrochloride | Dopamine | 2.22 | 208.61 |
| CP-471474 | Cytoskeleton and Extracellular Matrix | 2.22 | 208.56 |
| Brefeldin A from Penicillium brefeldianum | Cytoskeleton and ECM | 2.2 | 207.47 |
| Isoxanthopterin | Cell Stress | 2.2 | 207.3 |
| PD 0325901 | Kinase phosphotase biology | 2.18 | 206.39 |
| DL-Cycloserine | Sphingolipid | 2.17 | 206.2 |
| Indatraline hydrochloride | Dopamine | 2.17 | 205.97 |
| Podophyllotoxin | Cytoskeleton and ECM | 2.16 | 205.54 |
| (±)-Methoxyverapamil hydrochloride | Ca2+ Channel | 2.16 | 205.32 |
| Altretamine | DNA Metabolism | 2.15 | 205.19 |
| (±)-CGP-12177A hydrochloride | Adrenoceptor | 2.15 | 204.76 |
| N6-Cyclohexyladenosine | Adenosine | 2.12 | 203.65 |
| Lidocaine hydrochloride | Na+ Channel | 2.11 | 202.72 |
| (S)-Propranolol hydrochloride | Adrenoceptor | 2.1 | 202.63 |
| ML-7 | Phosphorylation | 2.09 | 201.69 |
| Ivermectin | Cholinergic | 2.07 | 201.04 |
| LE 300 | Dopamine | 2.06 | 200.3 |
| (±)-AMT hydrochloride | Nitric Oxide | 2.06 | 200.2 |
| Maraviroc | Immune signaling | 2.01 | 197.7 |

**Supplemental Table S2.** Target sequences for siNUDT5 #1-2

| siRNA Name | Target Sequence |
|---|---|
| siNUDT5 #1 | 5′-CATCGTGACAGTCACCATTAA-3 |
| siNUDT5 #2 | 5′-CAAGAACCAACGGAATCTTCT-3 |
| siControl | All Stars Negative Control siRNA (Qiagen) |

**Supplemental Table S3.** Primers used for qRT-PCR analysis.

| Gene Name | Forward Primer | Reverse Primer |
|---|---|---|
| NUDT5 v1 | CCC TAC ACC TTG GAG GTG AAC T | GAT TCC GTT GGT TCT TGG CTC |
| NUDT5 v2 | CCG AAG CCA AAG CCA GAG TT | TCA GCT ACC AGA GCA TCA AGT |
| NUDT5 v3 | AGA GAG AGC TAT GCC AAA AGA ACT | AAG TGT TCT CTG CAG CAC GG |
| PUM1 | CAG ACC AGC AGG TAA TTA ATG AGA | TTG CAA AGA CTG GGG CTG TA |

**Supplemental Table S4**. Antibody information for immunofluorescence and Western blotting analysis

| Antibody | Vendor | Catalog # | Dilution | Application |
|---|---|---|---|---|
| NUDT5 | Santa Cruz | sc-398644 | 1:25<br>1:50 | IF<br>WB |
| pFAK | Cell Signaling | 3283S | 1:1500 | WB |
| FAK | Cell Signaling | 71433S | 1:1500 | WB |
| pMLC2 | Cell Signaling | 3671S | 1:250<br>1:1500 | IF<br>WB |
| ppMLC2 | Cell Signaling | 95777S | 1:400<br>1:1500 | IF<br>WB |
| MLC2 | Sigma | M4401 | 1:1500 | WB |
| Vimentin | Cell Signaling | 5741S | 1:400 | IF |
| Myosin IIA | Cell Signaling | 49349S | 1:500 | IF |
| Myosin IIB | Cell Signaling | 8824S | 1:200 | IF |
| α-Tubulin | Cell Signaling | 2144S | 1:100 | IF |
| HSP90 | Santa Cruz | sc-13119 | 1:3000 | WB |

IF: Immunofluorescent staining; WB: Western blot

**Supplemental Table S5.** Adenosine aptamer and quencher sequences

| Name | Sequence |
|---|---|
| Active aptamer | FAM-CAC TGA CCT GGG GGA GTA TTG CGG AGG AAG GT |
| Inactive aptamer | FAM-GTG AAG GTG CAA CAG TCG GCT GGA GTT CGG AG |
| Active quencher | TCC CCC AGG TCA GTG/BHQ_1/ |
| Inactive quencher | CTG TTG CAC CTT CAC/3BHQ_1/ |
